# Lineage-tracing of *Acta2*+ cells in aged mice during lung fibrosis formation and resolution supports the lipofibroblast to myofibroblast reversible switch

**DOI:** 10.1101/2024.02.19.580952

**Authors:** Arun Lingampally, Marin Truchi, Olivier Mauduit, Vanessa Delcroix, Stefan Hadzic, Janine Koepke, Ana Ivonne Vazquez-Armendariz, Susanne Herold, Christos Samakovlis, Helen P Makarenkova, Elie El Agha, Chengshui Chen, Bernard Mari, Saverio Bellusci

## Abstract

Idiopathic pulmonary fibrosis (IPF) develops mostly in old man and is characterized by the irreversible accumulation of excessive extracellular matrix components by activated myofibroblasts (aMYFs) leading to lung failure. Following bleomycin administration in young mice, fibrosis formation associated with efficient resolution takes place, the later limiting the clinical relevance of this model for IPF. In young mice, we previously reported that aMYFs captured during fibrosis formation differentiate towards a lipofibroblast (LIF)-like phenotype during resolution.

In this study, we used aged mice in combination with bleomycin administration to trigger enhanced fibrosis formation and delayed resolution, as a more relevant model for IPF and examined the heterogeneity and fate of aMYFs at different time points. Alveolosphere assay were carried out to compare the alveolar resident mesenchymal niche activity for AT2 stem cells in young versus old mice. Lineage tracing of the Acta2+ aMYFs in old mice exposed to bleomycin followed by scRNAseq of the lineage-traced cells isolated during fibrosis formation and resolution was performed to delineate the heterogeneity of aMYFs during fibrosis formation and their fate during resolution. Data mining of human mesenchymal cells from IPF and control datasets were also performed to decipher the heterogeneity of aMYFs and investigate differentiation trajectories during fibrosis formation.

Our results show that alveolar resident mesenchymal cells from old mice display decreased supporting activity for AT2 stem cells. We report that aMYFs consist of four subclusters displaying unique pro-alveologenic versus pro-fibrotic profiles. Alveolar fibroblasts displaying a high LIF-like signature largely constitute both the origin and fate of aMYFs during fibrosis formation and resolution, respectively. The heterogeneity of aMYFs is conserved in humans and a significant proportion of human aMYFs displays a high LIF signature.

In conclusion, our data indicate that the cellular and molecular bases of aMYFs formation and differentiation towards the LIF phenotype are conserved between young and old mice. Importantly, our work identifies a subcluster of aMYFs that is potentially relevant for future management of IPF.

## Introduction

Idiopathic pulmonary fibrosis (IPF) is characterized by the accumulation of excessive extracellular matrix (ECM) components due to an aberrant wound-healing process. The initiation of the IPF is widely proposed to be starting in the alveolar epithelium (Selman et al., 2001; Selman & Pardo, 2006), either due to repetitive injury or aging. Chronic alveolar epithelium injury in the context of aging is proposed to lead to the activation of the adjacent mesenchyme, progressively resulting in the formation of the activated myofibroblasts (aMYFs).

The bleomycin (bleo) model is often used to study fibrosis in young mice. However, it is criticized as fibrosis formation is also associated with efficient resolution. In contrast, IPF in humans is a progressive disease which fibrosis never resolves. To overcome this caveat linked to the use of a single bleo administration to young mice, alternative approaches such as repetitive-bleo injury in young mice have been proposed. Three successive weekly doses of bleo in young mice trigger increased fibrosis formation at 2 weeks. Subsequent fibrosis resolution, which is thought to be finalized at 4 weeks after a single bleo dose, is instead finalized at 6 weeks after the final dose (Chung et al., 2003). In addition, another study using eight biweekly bleo doses showed persistent fibrosis up to 10 weeks after the final dose (Degryse et al., 2010).

Another alternative approach to reflect human IPF is to use aged mice, which were reported to develop severe fibrosis with impaired resolution (Caporarello et al., 2020; Redente et al., 2011). Aging is associated with telomere shortening, cellular senescence, oxidative stress, and stem cell exhaustion (Bueno, Calyeca, Rojas, & Mora, 2020; Chanda et al., 2021; Cronkhite et al., 2008; Sgalla et al., 2018; Yanai et al., 2015). A recent study showed that the niche activity of the resident alveolar mesenchymal cells (rMCs) for the AT2 stem cells is lost within the aged mesenchyme due to increased expression of NADPH oxidase 4 (Nox4), an enzyme essential for regulating reactive oxygen species. The rMC-AT2 stem cell niche activity is partially rescued with the partial loss of Nox4 in aged rMCs (Chanda et al., 2021). In addition, NOX4 is elevated in IPF lungs. Interestingly, siRNA-based inhibition of NOX4 led to a decrease in SMAD2/3 phosphorylation which is downstream of Transforming growth factor beta (TGFβ) signaling, a master signaling cascade in IPF (Amara et al., 2010).

As aforementioned, the initiation of the IPF is proposed to start in the alveolar epithelium. Focusing on the epithelium, a recent study showed the effects of aging and injury on the alveolar epithelial type 2 (AT2) cells. This study observed that AT2s are significantly decreased in aged mice and that new AT2 subsets emerge, which are enriched in the expression of inflammation, senescence, and apoptosis markers. Additionally, young AT2s increased the aging-related markers and shifted towards these new AT2 subsets after bleo injury. However, young AT2s recovered back to their native state during fibrosis resolution. In contrast, aged AT2s failed to recover back during fibrosis resolution. This suggests that aging impairs the AT2s progenitor renewal capability (Liang et al., 2023).

Considering the AT2s role as progenitors, the Cell division control protein 42 homolog (Cdc-42) is essential for the differentiation of AT2 cells to alveolar epithelial type 1 (AT1) cells. In mice, the deletion of *Cdc-42* in AT2 cells at a young age triggered fibrosis formation from the periphery to the center of the lung with aging (Wu et al., 2020). These studies show that aged mice are primed for fibrosis compared to young mice. Recently, studies comparing young and old mice revealed that pathways associated with fibrosis are similarly regulated between young and aged mice. Aged mice, however, showed an increase in immune cells. Additionally aged mice showed delayed fibrosis resolution which could be due to decreased regenerative capacity of the aged lung (Klee et al., 2023; Weckerle et al., 2022).

Our previous studies showed that lipofibroblasts (LIFs), defined as resident mesenchymal cells located in the alveoli at proximity of alveolar type 2 epithelial cells (O’Hare & Sheridan, 1970; Vaccaro & Brody, 1978) are essential for the proliferation of AT2 stem cells (Taghizadeh et al., 2022; Taghizadeh et al., 2021). LIFs are also a potential source of aMYFs. Lineage tracing of LIFs and Acta2+MYFs in the context of fibrosis formation and resolution in young mice demonstrated a LIF to MYF reversible switch (El Agha et al., 2017). Recently, using lineage tracing of the Acta2+MYFs associated with scRNASeq, we reported the cellular and molecular cues associated with the LIF-to-aMYF reversible switch during fibrosis formation and resolution in young mice using the bleo model (<u>Lingampally et al., 2024 in revision at ERJ</u>). We found that the aMYFs are heterogeneous and express *Cthrc1*. Our data suggest that LIFs differentiate to Cthrc1+aMYFs during fibrosis formation and switch back to their native LIF phenotype during resolution.

In contrast, whether a LIF-to-aMYF reversible switch takes place in aged mice is still unclear. To explore the LIF-to-aMYF reversible during aging, we employed the 52-56 weeks old *Tg(Acta2-CreERT2)/+; tdTom^flox^*mice, which replicates the mid-age of humans where initiation of the fibrosis formation in proposed to take place (Jenkins et al., 2017).

In this study, alveolosphere assay were carried out to compare the alveolar resident mesenchymal niche activity for AT2 stem cells in young versus old mice. Lineage tracing of the Acta2+aMYFs in old mice exposed to bleo followed by scRNAseq of the lineage-traced cells isolated during fibrosis formation and resolution was performed to delineate the heterogeneity of aMYFs during fibrosis formation and their fate during resolution. Data mining of human IPF datasets was also performed to decipher the heterogeneity of aMYFs and investigate differentiation trajectories during fibrosis formation.

Our data indicate that the cellular and molecular bases of aMYFs formation and differentiation towards the LIF phenotype are conserved between young and old mice. Importantly, our work identifies a subcluster of aMYFs that is potentially relevant for future management of IPF.

## Materials and Methods

### Ethical aspects and Mice

All animal experiments were performed according to the approved protocols by the Regierungspraesidium Giessen, the animal ethics committee of the University of Giessen (permit numbers: G57/2019–No.974_GP and G26/2020–No.1002_GP). All animals were housed under specific pathogen-free (SPF) conditions with unlimited access to food and water. The room environment was maintained with 12 hours of dark/light cycle at 22°C and 40-70% humidity. *Tg(Acta2-CreERT2)* mice (STOCK_*Tg(Acta2-cre/ERT2)12Pcn*) (kind gift from Dr. Pierre Chambon, University of Strasbourg, France) were bred with *td-Tomato^flox^*(*B6;129S6-Gt (ROSA) 26Sortm9(CAG-tdTomato)Hze/J* (Jackson lab, 007909) to generate tamoxifen mediated reporter mice.

### Bleo instillation

52-56 weeks-aged female mice were intratracheally (IT) subjected to either saline or bleo 2 U/kg body weight (2 mg/kg) (Bleomedac, PNZ-02411351). Lungs were harvested at days 14, 30 and 60 post intratracheal instillation for analysis.

### Tamoxifen administration

Tamoxifen (Sigma, T5648-5G) was reconstituted in corn oil. Mice were intraperitoneally (IP) subjected (0.1 mg/g body weight) with four injections on days 5, 7, 9 and 11 *after* saline or bleo instillation, respectively.

### Immunofluorescence staining

Mice were sacrificed and the lungs were transcardially perfused with 10 mL PBS, perfused lungs were fixed in 4% PFA and sequentially dehydrated with ethanol. The fixed-dehydrated lungs were then embedded using paraffin. Five-μm-thick paraffin slices were deparaffinized and rehydrated, sequentially with xylol, 100%, 70%, 50% and 30% ethanol and finally in Milli-Q water. The slices were then subjected to antigen retrieval, by transferring the slices in citrate buffer and boiled for 15 min in a cooker. Following antigen retrieval, slices were cooled on ice for 20 min and washed with PBST three times. The slices were blocked with 3% BSA + 0.2% Triton-X in PBS for 1 h. Slices were then incubated with primary antibodies: anti-Acta2 (Sigma, F3777), anti-RFP (Invitrogen, R10367) at 4°C overnight. Following washing, slices were incubated with secondary antibodies for 1hour at room temperature. Finally mounted with ProLong Gold Antifade Reagent containing DAPI (Molecular Probes, P36935).

### Hematoxylin and eosin staining

Five-μm-thick paraffin slices were deparaffinized and rehydrated. The sections were stained with hematoxylin (Roth, T865.2) for 2 min and eosin (Thermo Fisher Scientific, 6766007) for 1 min. Following staining slices were dried and mounted.

### FACS preparation

Mice were sacrificed and the lungs were transcardially perfused with 10 mL PBS. Lungs were finely chopped and digested in 0.5% collagenase type IV (Gibco 17104-019) at 37°C for 45 min. Following incubation, the cell suspension is passed through 70 and 40 µm cell strainers. The single-cell suspension was centrifuged, and the cell pellet was stained with anti-Cd31 (Alexa Fluor 488-conjugated 1:100, Biolegend, 102514), anti- Cd45 (Alexa Fluor 488-conjugated 1:100, Biolegend, 103122), anti-Epcam (APC- Cy7-conjugated 1:50, Biolegend, 118217) and anti-Sca1 (pacific blue-conjugated 1:50, Biolegend, 108120) antibodies on ice for 20 min. After washing, SYTOX^TM^-Blue (Invitrogen, S34857) was added to cell suspension to exclude dead cells and sorting was carried out using FACSAria III cell sorter (BD Biosciences). Data were analyzed using FlowJo software (FlowJo, LLC).

### Generation of the scRNA-seq data

Live sorted tdTom+ cells were centrifuged and resuspend in 0.04% ultrapure BSA (Invitrogen, 01266574) in PBS for optimal cell concentration. The resuspended tdTom+ cells were loaded into the Chromium Controller (10x Genomics) and the cDNA libraries were prepared according to manufacturer’s instructions. Sequencing was performed by Nextseq2000 (Illumina, Inc.) and reads were aligned against to a custom mouse reference genome (mm10) and counted by STARsolo.

### Analysis of the scRNA-seq data

All downstream analyses were carried out with Seurat R package (v4.1.0) (Y. Hao et al., 2021). In order to be consistent with the cells annotation produced for the young Acta2+ mice dataset (<u>Lingampally et al., 2024 in revision at ERJ</u>), the four old mice samples (saline d14, Bleo-Tam d14, Bleo-Tam d30 and Bleo-Tam d60) were integrated with 3 young mice samples (saline d14, Bleo-Tam d14 and Bleo-Tam d60). First, the counts matrix of each sequenced sample were loaded as seurat objects, then filtered using arbitrary thresholds for UMI, genes and mitochondrial content (nCount_RNA < 30000, nFeature_RNA > 800 and percent.mito < 0.15), before to be concatenated. The concatenated matrix was normalized using the NormalizeData function. After normalization, the counts of the 2000 most variable features selected with “vst” method were scaled and centered before running PCA. Harmony (Korsunsky et al., 2019) was run on the PCA cell embedding, specifying the sample of origin as covariate. kNN clustering was run on the first 50 dimensions computed with Harmony (using k.param =10), same for UMAP visualizations in two-dimensions. Based on UMI content and differentially expressed genes in each cluster, clusters corresponding either to remaining low quality cells or non-mesenchymal cells were removed. The integration process was run again on the subset of selected cells with the same parameters. Finally, cell clusters were annotated according to the expression of their specific markers described in (Tsukui et al., 2020). Subpopulations of *Cthrc1+* aMYFs, alveolar (LIF), adventitial and peribronchial fibroblasts were identified by performing clustering on each particular cell subsets, using the same first 50 dimensions computed with Harmony on the complete dataset. Differential expression between samples for a particular subpopulation were run using the FindMarkers function on normalized data.

### Data deposition

Single cell data have been deposited on gene expression omnibus (GEO) repository: GSE253453, GSE221402 and GSE223664.

### Human scRNAseq data mining

To analyze the stromal compartment of human lungs, we took advantage of the recently published human lung cell atlas (HLCA) core dataset (Sikkema et al., 2023) available on CELLxGENE Census (https://cellxgene.cziscience.com) and combined it with samples from the following publicly available IPF datasets: GSE136831 (Adams et al., 2020), GSE227136 (Natri et al., 2023), GSE132771 (Tsukui et al., 2020), GSE128033 (Morse et al., 2019). For our analysis, only samples from lung parenchyma of non-cancer patients were retained from the HLCA core dataset. From the HLCA, GSE136831 and GSE227136 datasets, we selected all cells annotated as ‘stromal’ by authors. For GSE132771 and GSE128033, we performed unsupervised clustering independently and used Azimuth (https://azimuth.hubmapconsortium.org) for label transfer from HLCA annotations. We thus kept for downstream analysis all cells with a prediction score > 0.5 for the following cell identities (predicted annotation level 2): ‘fibroblast lineage’, ‘mesothelium’, ‘smooth muscle’. For all datasets, cells with more than 10% mtDNA and samples containing less than 100 stromal cells were excluded. The resulting list of samples included 62284 cells in total (26391 control and 35893 IPF cells). Samples were normalized using SC Transform v2 and integrated with reciprocal PCA using R/Seurat (v5, (Yuhan Hao et al., 2022)). Unsupervised clustering and UMAP reduction allowed the annotation of the stromal subtypes. The statistical analysis of cell proportions in IPF and control samples was conducted with the R package speckle (Phipson et al., 2022).

For the trajectory analysis on the fibroblast subset, we used slingshot (Street et al., 2018) with the dimensionality reduction produced by PCA and cluster labels previously identified. Then, we subsetted cells from the inferred trajectory that belonged to the LIF, alveolar FIB and aMYFs clusters and performed a similar analysis by setting aMYF as the end cluster. For visualization purposes, the resulting trajectories, lineages and principal curves were projected onto the UMAP reduction plot. To identify temporally dynamic genes and genes differentially expressed between control and IPF samples along the trajectory, we used tradeSeq (Van den Berge et al., 2020) that fits a generalized additive model (GAM) to smooth each gene’s expression depending on the condition (control or IPF). Smoothed expression estimates were computed with 5 knot points for the common 6413 variable genes we independently identified in control and IPF datasets. Gene overlap and pathway analysis were conducted with Metascape (metascape.org) by interrogating GeneOntology, WikiPathways and Reactome databases. For all comparisons, genes significantly associated with pseudotime or differentially expressed were identified using the Wald test with a log2 fold-change cut- off at +/- log2(2) and FDR < 0.05.

### Alveolosphere assay

Alveolosphere assay was carried out as previously described (Taghizadeh et al., 2021). The following mice were used to sort resident mesenchymal cells (rMCs) and Alveolar type 2 cells (AT2s): 1) rMCs from C57BL/6 mice were isolated from 8 weeks-old (young) and 35 weeks-old (mature) mice. rMCs from *Tg(Acta2-CreERT2)/+; tdTom^flox^*mice were isolated from 52-56 weeks-old (aged) mice. 2) AT2s from *S*ftpc*^CreERT2/+^; tdTom^flox^* mice were isolated from 35 weeks-old mice. AT2s from C57BL/6 mice were isolated from 8 weeks-old mice.

In brief, sorted 50,000 rMCs and 5,000 AT2s were reconstituted in 100 µL media, and mixed in of 100 µL ice cold Matrigel (Corning, 356231). The mixture was transferred to 24-well 0.4 µm transwell inserts (Greiner bio-one, 662641) and incubated at 37°C for 15 min for Matrigel polymerization. Next, 500 µL of media (sorting media plus 1% ITS (Gibco, 41400-045)) was added to each well and incubated at 37°C in 5% CO_2_ for 14 days.

### Statistical analysis

For comparison of two groups, unpaired, two-tailed t-tests were used. For comparisons involving more than two groups, one-way ANOVA with post hoc Newman–Keuls multiple comparisons test or 2-way ANOVA with post hoc Šídák’s multiple-comparisons test was used. Values of P<0.05 were considered statistically significant. All data are presented as mean ± SEM.

## Results

### Functional characterization of the rMC niche activity for AT2 stem cells in young and aged mice

The alveolosphere assay was carried out to determine the effect of aging on the rMC niche activity for AT2 stem cells. To generate the organoids, rMCs Sca1+ from young (8 weeks-old) and mature (35 weeks-old) mice were isolated by FACS. rMCs Sca1+ are enriched in LIFs displaying the niche activity for AT2 stem cells (McQualter et al., 2013; Taghizadeh et al., 2022; Taghizadeh et al., 2021).

rMCs Sca1+ were co-cultured with tdTom^High^-AT2s from 35 weeks-old *S*ftpc*^CreERT2/+^; tdTom^flox^*mice. The gating strategy to sort rMCs Sca1+ and tdTom^High^ AT2s is shown in Figure 1a. While young rMCs Sca1+ co-cultured with mature AT2s in Matrigel for 14 days. led to the formation of organoids with the expected CFE and size (Taghizadeh et al., 2021), rMCs Sca1+ from 35 weeks-old mice failed to display such niche activity (Figure 1b,c) Given that the 35 weeks-old mesenchyme lost the niche activity, which we propose to be due to a progressive transition of the LIFs towards a MYF-like phenotype, we decided to use 52-56 weeks-old (aged) mice to investigate the process of fibrosis formation and resolution. 52-56 weeks-old *Tg(Acta2-CreERT2)/+; tdTom^flox^* female mice were intratracheally (IT) administered with saline (control mice) or bleomycin (bleo) at a dose which was reported to induce moderate fibrosis in young female mice. Tamoxifen was intraperitoneally injected (Tam-IP) on days 5, 7, 9 and 11 to label the *Acta2*+ cells. Next, we sorted rMCs Sca1+ from saline and bleo mice at day 14 using the gating strategy described in Figure 1d. AT2s from 8 weeks-old C57BL/6 mice were sorted as described in Figure 1d and co-cultured with aged saline and bleo rMCs Sca1+ in Matrigel for 14 days. Our results indicate that organoid formation is completely lost in aged saline mice (Figure 1e) confirming that the underlying cause for this result is the loss of the niche activity in the mesenchyme and not deficient AT2s, which in our conditions are coming from young mice and are therefore displaying robust AT2 stem cells. As previously described, bleo injury also causes loss of the niche activity (Figure 1e,f).

**Figure 1:**
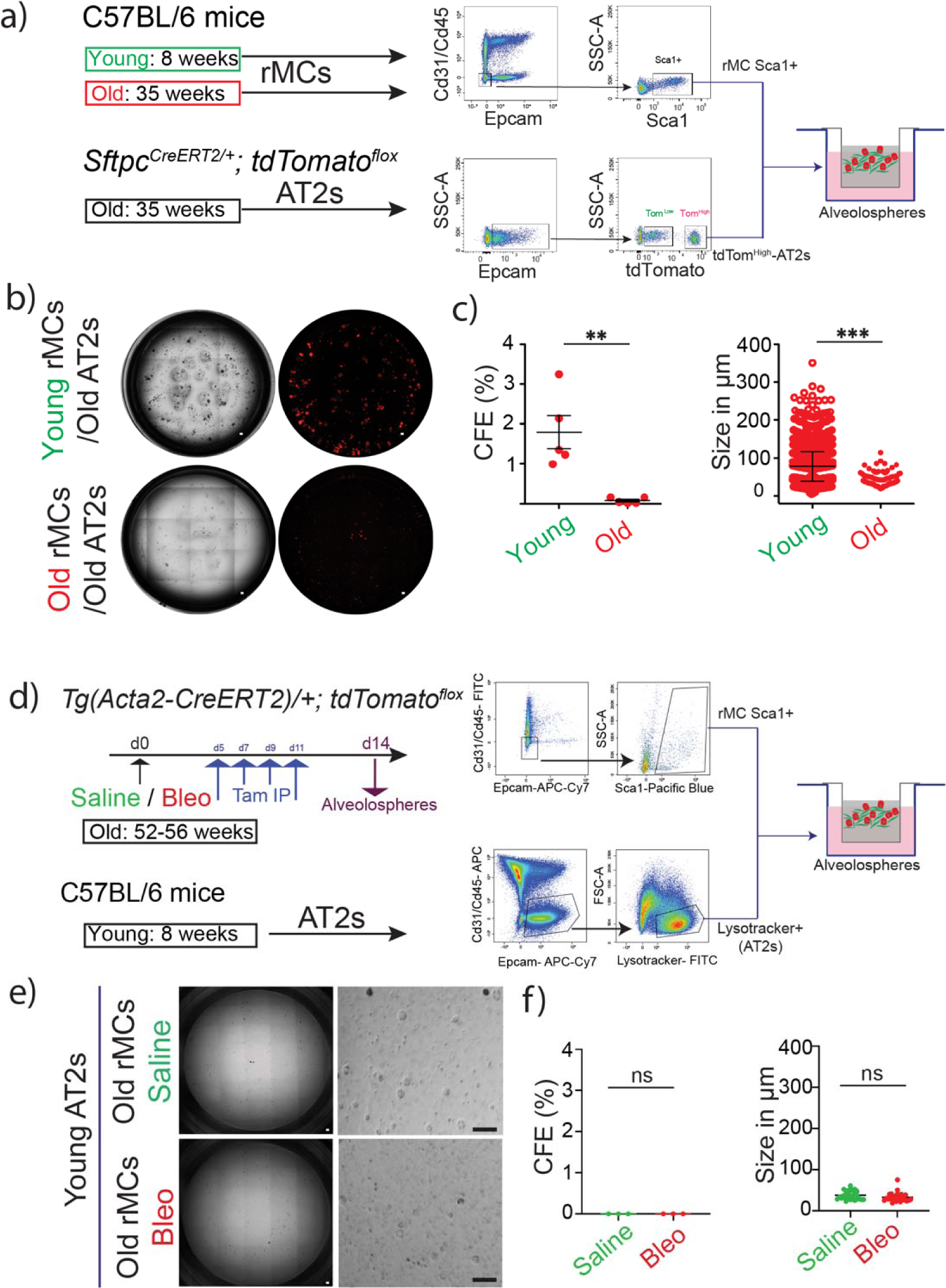
E**valuation of resident mesenchymal niche activity for AT2 stem cells upon aging. a)** Experimental design showing the rMCs (resident mesenchymal cells) isolated either from 8 or 35 weeks C57BL/6 mice and gating strategy to sort Cd31/Cd45/Epcam triple negative (rMC) positive for Sca1 (rMC Sca1+ cells). Experimental design showing the AT2 cells (Alveolar type 2 cells) isolated from 35 weeks SftpcCreERT2)*^/+^;tdTomato^flox^* mice and gating strategy to sort Epcam+ and tdTomato+ (Tom^High^) cells. AT2 and rMC Sca1+ cells are co-cultured in Matrigel for 14 days (alveolosphere assay). **b)** Bright field and fluorescent images showing formation of organoids from old-AT2s/Tom^High^ (35 weeks mice) cells co-cultured with rMC Sca1+ cells are isolated from Young-C57BL/6 (8 weeks) mice and such resident mesenchymal niche activity is lost in Old-C57BL/6 (35 weeks) mice. **c)** Quantification of colony forming efficiency confirming the loss in stem cell niche activity upon resident mesenchymal aging (n=3/condition). **d)** Experimental design showing the rMCs isolated either from saline or Bleomycin injured mice at day 14 post-bleo injury from Old-*Tg(Acta2- CreERT2)/+;tdTomato^flox^* (52-56 weeks) female mice, and gating strategy to sort Cd31/Cd45/Epcam triple negative (rMC) positive for Sca1 (rMC Sca1+ cells). Experimental design showing the AT2 cells isolated from non-injured Young-C57BL/6 (8 weeks) mice, and gating strategy to sort Epcam+ and Lysotracker+ (AT2) cells. AT2 and rMC Sca1+ cells are co-cultured in Matrigel for 14 days (alveolosphere assay). **e)** Bright field images showing organoids failed to form from saline or bleo-injured mice (few cysts are formed in both conditions, which are around 50 µm in size). **f)** Quantification of colony forming efficiency showing the loss in stem cell niche activity in saline or bleo-injured mice upon resident mesenchymal aging and no significant changes is seen in the size (n=3/condition). Scale bar: b, e- 100 µm. Statistical analysis was performed using: (c, f)- Unpaired two-tailed t-test. **: P<0.01; ***: P<0.001

To summarize, these results indicate that in the alveolosphere model and in our experimental conditions, mature AT2s are not the limiting factor for organoid formation, However, the niche activity displayed by rMCs Sca1+ is significantly impaired with aging.

### Lineage tracing of *Acta2+* cells in aged healthy and fibrotic mouse lungs

52-56 weeks-old *Tg(Acta2-CreERT2)/+; tdTom^flox^* female mice were intratracheally (IT) administered with saline (control mice) or bleo with 2 U/kg body weight, a dose which was reported to induce moderate fibrosis in young female mice (Lingampally et al., 2024 in revision at ERJ). Mice were injected with Tam-IP post saline or after bleo administration to label *Acta2+* cells and examine their heterogeneity during fibrosis formation at d14 and their fate during resolution at d30 and 60 following bleo administration (Figure 2a). Experimental (bleo) and control (saline) mice were first examined on day 14, at the peak of fibrosis. Hematoxylin Eosin staining showed abundant fibrotic areas upon bleo administration (Figure 2b).

**Figure 2:**
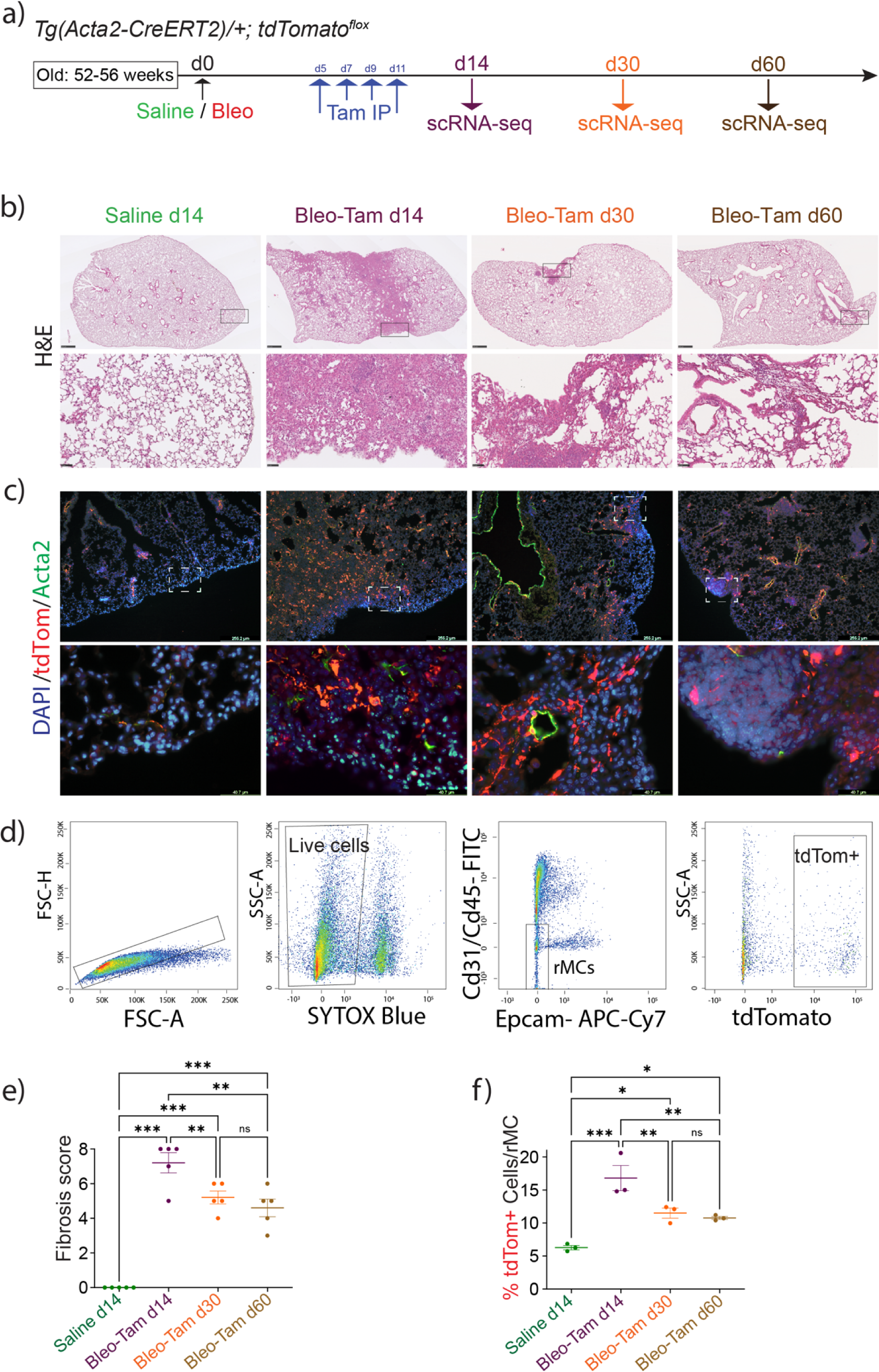
Lineage tracing of *Acta2*+ cells in saline and bleomycin injured mice. a) 52-56 weeks-old female *Tg(Acta2-CreERT2)/+;tdTomato^flox/flox^* mice are used to lineage label *Acta2*+ cells in saline, and bleo-injured mice. Control lungs are collected at day14 following saline administration and experimental lungs (Bleo-Tam) are collected at d14, d30 and d60 following bleo injury. **b)** Corresponding low and high magnification of H&E staining showing fibrosis formation at Bleo-d14 peak of fibrosis and fibrotic lesions/regions are still present at resolution Bleo-d30 and Bleo-d60 following bleomycin injury. **c)** Corresponding low and high magnification of IF staining against tdTom, Acta2 and DAPI, showing the presence of tdTom+ cells in the bronchiolar region as well as in the alveolar region in saline conditions. However, abundant recruitment of tdTom+ cells in the fibrotic regions in Bleo-d14, Bleo-d30 and Bleo-d60 lungs following bleo injury. **d)** Gating strategy to sort SYTOX- negative live lineage-labeled tdTom+ cells. **e)** Ashcroft score confirming massive formation in old mice upon bleo injury (b), saline d14 0 ± 0 score, Bleo-Tam d14 7.2 ± 0.6 score, Bleo-Tam d30 5.2 ± 0.4 score and Bleo-Tam d60 4.6 ± 0.5 score. **f)** Quantification of sorted tdTom+ cells over rMC (d) from saline d14 6.3% ± 0.3, Bleo-Tam d14 16.8% ± 1.9, Bleo-Tam d30 11.5% ± 0.8 and Bleo-Tam d60 10.8% ± 0.2. Scale bar: b: Low magnification- 500 µm, High magnification- 50 µm; c: Low magnification- 256.2 µm, High magnification- 40.7 µm. Statistical analysis was performed using: (e and f)- One-way ANOVA with Newman–Keuls post hoc test for multiple comparisons. *: P<0.05; **: P<0.01; ***: P<0.001.

Furthermore, Bleo-Tam d30 and Bleo-Tam d60 lungs demonstrated the presence of significant residual fibrotic areas in the lung parenchyma (Figure 2b). Ashcroft scoring (Ashcroft, Simpson, & Timbrell, 1988) at d14 was significantly increased in experimental compared to control lungs confirming massive fibrosis formation. Ashcroft scoring at day 30 and 60 indicated only partial fibrosis resolution (Figure 2e).

Next, we examined if aged mice compared to young mice (<u>Lingampally et al., 2024 in</u> <u>revision at ERJ</u>) displayed, for the same bleo dose, increased fibrosis and delayed resolution by comparing their respective Ashcroft score. As expected, old mice displayed increased Ashcroft score both at d14 and at d60 (Figure S1b,c) indicating elevated fibrosis formation and delayed fibrosis resolution compared to young mice as previously described.

Next, we examined the lineage-traced *Acta2+* (tdTom*+*) cells by IF using antibodies against RFP and alpha-smooth muscle actin (Acta2). In saline conditions, we found tdTom+ cells located within the airway and vascular smooth muscle cells (ASMCs and VSMCs) co-expressing high levels of Acta2 (Figure 2c). Additionally, we also found tdTom+ cells with low levels of Acta2 in the parenchyma of the lung (Figure 2c). Next, we examined tdTom+ cells at the peak of fibrosis (day 14) in bleo-treated mice. We observed massive recruitment of tdTom+ cells in the fibrotic regions. Interestingly, tdTom+ cells are still abundantly present in the lung both at d30 and d60 (Figure 2c). Flow cytometry was used to sort and quantify the percentage of tdTom+ cells out of the Cd45/Cd31/Epcam triple-negative resident mesenchymal cells (rMCs) in saline d14, Bleo-Tam d14, Bleo-Tam d30 and Bleo-Tam d60 lungs. Representative FACS plots are shown in Figure 2d. SYTOX^TM^ staining was used to exclude dead cells from the analysis and to sort live tdTom+ cells for scRNA-seq experiments. FACS-analysis confirmed a very significant increase of the percentage of tdTom+ cells over rMC in Bleo-Tam d14 (16.8% ± 1.9) compared to saline d14 (6.3% ± 0.3). We also detected the significance presence of tdTom+ cells at Bleo-Tam d30 (11.5% ± 0.8) and Bleo-Tam d60 (10.8%± 0.2) compared to saline d14 (Figure 2f).

### *Acta2*+ cells lineage-labelled during fibrosis formation massively contribute to the *Cthrc1*+ aMYF lineage

Lineage-labeled *Acta2*+ (tdTom+) cells were sorted from saline d14 and Bleo-Tam lungs at d14, d30 and d60 by negative gating for Cd31, Cd45 and Epcam (Figure 2d). Then, 9,000 cells per condition were loaded for scRNA-seq. After quality control (Figure S2, Table 1), a total of 25714 cells were recovered for the integrated analysis, 9088 cells from the saline d14, 3239 cells from Bleo-Tam d14, 5700 cells from Bleo-Tam d30 and 7687 cells from Bleo-Tam d60. The integrated UMAP showed that tdTom+ cells contribute to 16 different subclusters including alveolar fibroblasts, adventitial fibroblasts (1,2), peribronchial fibroblasts (1-3), inflammatory fibroblasts, smooth muscle cells (SMC)/pericytes, proliferating and *Cthrc1*+ fibroblasts (Ct1-4) (Figure 3a and b). Figure 3c shows the heatmap for the genes enriched for each cluster. The *Cthrc1+* fibroblasts are enriched with fibrotic genes such as *Spp1*, *Ltbp2* and *Col1a1*.

**Figure 3:**
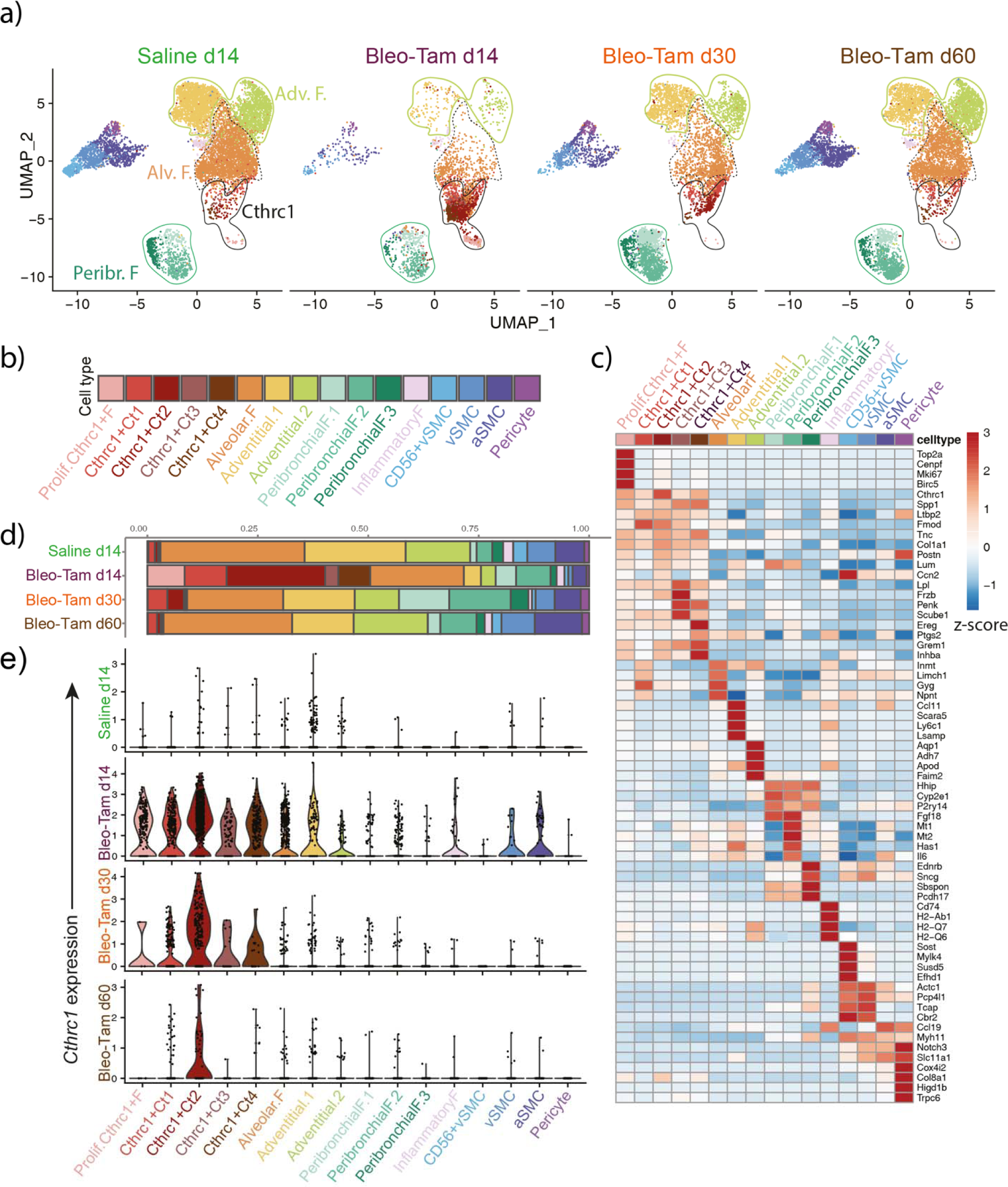
A*c*ta2*+* cells massively contribute to the *Cthrc1+* myofibroblast lineage during fibrosis. a) Integrated UMAP of *Acta2+* cells isolated from saline d14 and Bleo- Tam d14, d30 and d60 lungs, showing 16 distinct clusters including Alveolar fibroblasts, peribronchial fibroblasts, adventitial fibroblasts and *Cthrc1+* myofibroblasts. b) Nomenclature/ cell identity of each cluster. c) Heatmap of subpopulations markers. d) Relative frequencies of each subpopulation in saline d14 and Bleo-Tam d14, d30 and d60 lungs. Note that the *Cthrc1+* myofibroblasts are massively recruited (around 50%) in Bleo-Tam d14 lungs and proliferating *Cthrc1+* cells are also increased at peak of fibrosis, suggesting aging is a predisposition for lung fibrosis. e) Normalized expression of *Cthrc1* in the different subpopulations in saline d14 and Bleo-Tam d14, d30 and d60 lungs. Note that the *Cthrc1* expression is high at day 14 peak of fibrosis, expression was maintained at day 30 and decreased during fibrosis resolution in day 60 lungs.

The relative frequencies of the different clusters at each time point is shown in Figure 3d. The distribution of clusters supports our previous observations made in young mice (Lingampally et al., 2024 in revision at ERJ). We observe that *Cthrc1+* myofibroblasts are increased upon bleo injury. Proliferating and *Cthrc1+* myofibroblasts (Ct1-4) at day 14 massively increase to represent around 50% of the tdTom+ cells analyzed in bleo versus only 5% in saline d14.

The percentage of alveolar fibroblasts was also significantly decreased at d14 compared to saline. In contrast, at Bleo-Tam d30, during the early resolution phase, the proliferating and *Cthrc1*+ fibroblasts (Ct1-4) were decreased (around 10%). The relative percentage of alveolar fibroblasts was not changed. By the time of late resolution, at Bleo-Tam d60, the proliferating and *Cthrc1*+ fibroblasts (Ct1-4) reached the basal level observed in saline level (around 5%). The alveolar fibroblasts showed an increased in their percentage at that time point. Violin plots indicated that proliferating and all *Cthrc1+* fibroblast subclusters (Ct1-4) express high levels of *Cthrc1* with Ct2 showing the maximum expression. Indeed high levels of *Cthrc1* expression was observed at peak of fibrosis Bleo-Tam d14 and the expression was maintained at Bleo-Tam d30. Decreased *Cthrc1* expression in Bleo-Tam d60 was observed during late resolution (Figure 3e). To summarize, these results indicate that aged mice are more susceptible to fibrosis illustrated by an abundant *Cthrc1*+ population. In addition, we observed the maintenance of *Cthrc1* expression in Ct2 cells during fibrosis resolution at day 60.

### Deconvolution of the *Cthrc1+* myofibroblast cells heterogeneity and their fate during fibrosis formation and resolution

Next, we carried out a detailed analysis for the *Cthrc1*+ clusters from saline d14, Bleo- Tam d14, d30 and d60 lungs (Figure 4). All *Cthrc1*+ clusters were amplified in the context of fibrosis formation at d14. Indeed, the Ct2 (22.32%) cluster is most abundant over the other *Cthrc1*+ clusters such as Ct1 (9.51%), Ct3 (2.96%) and Ct4 (7.22%). However, at d30 and d60, during resolution, the relative percentages of *Cthrc1*+ (Ct1-4) clusters were significantly decreased (Figure 4a). Note that *Cthrc1+* myofibroblasts (Ct1-4) at day 14 massively increased from 25% in young mice (Lingampally et al., 2024) to 50% in old mice (Figure S3b and d). Interestingly, Ct2 is the most abundant cluster in both conditions, which shows the pivotal role of this population in fibrosis development.

**Figure 4:**
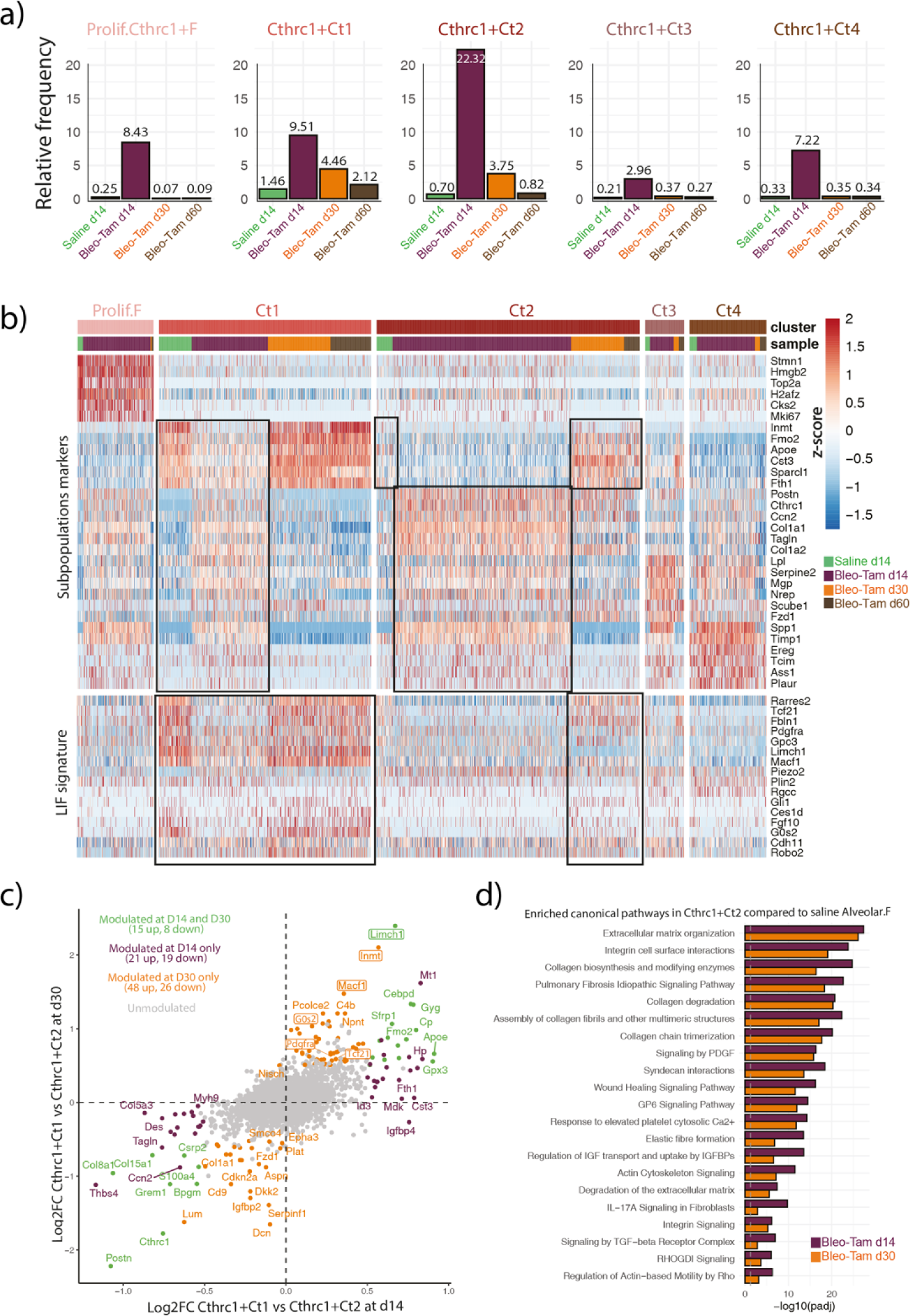
Comparison of *Cthrc1+* myofibroblast cells during fibrosis formation and resolution. a) *Acta2+* cells isolated from saline d14 and Bleo-Tam d14, d30 and d60 lungs contributing to 5 clusters (Prolif.F, Ct1, Ct2, Ct3 and Ct4) of *Cthrc1+* cells. Relative frequencies of each Cthrc1-expressing subpopulations at each time-point. Note that the *Cthrc1+* myofibroblasts clusters frequencies are high in Bleo-Tam d14 lungs. b) Heatmap showing the scaled expression of Cthrc1-expressing subpopulations markers (top) and genes of the lipofibroblast (LIF) signature (bottom) in saline d14 and Bleo-Tam d14, d30 and d60 lungs. **c)** Correlation of the two Log_2_ Fold change obtained by comparing *Cthrc1+* Ct1 to Ct2 at day 14 and at day 30 post bleo-injury. Green dots correspond to genes significantly modulated (|log_2_FC|>0.5 and padj < 0.05) in both time points. Purple and orange dots correspond respectively to genes only significantly modulated at day 14 or day 30. Highlighted genes are canonical markers of LIF. **d)** IPA® Canonical Pathways enrichment obtained from the differential analyses of Ct2 at day 14 or day 30 compared to saline d14 alveolar fibroblasts.

Figure 4b shows the heatmap of the most expressed gene markers for each *Cthrc1+* sub cluster. Ct1 contains *Inmt* and *Apoe*, two canonical Alveolar fibroblast markers. The Ct1 cluster is significantly impacted, which leads to a decrease in its signature following bleo injury. At the peak of fibrosis, at d14, this cluster adopts an intermediate Ct2, Ct3 and Ct4 signature (shown in black box). Interestingly, during early (d30) and late resolution (d60), the Ct1 signature retrieves back to its native saline state, with a slight increase at d60. The Ct2 cluster in saline displayed a Ct1 signature. However, upon bleo injury, the Ct2 cluster loses the Ct1 signature and differentiates to a Ct2 status which is enriched with fibrotic genes such as *Postn*, *Col1a1*, *Cthrc1* and *Tagln*. Additionally, this cluster also adopts an intermediate Ct3 and Ct4 signature (shown in black box). Interestingly, during early (d30) and late (d60) resolution, the Ct2 signature retrieves back to its native saline/Ct1 status. Both Ct4 and Ct3 showed a similar trend like Ct1 and Ct2 during fibrosis formation at d14 and resolution at d60. However, due to the low number of cells during resolution, the shift toward the Ct1 signature is not clearly seen.

Next, when we plugged-in the previously described LIF signature into *Cthrc1+* (Ct1-4) clusters during fibrosis formation and resolution, Ct1 shows the highest level of all clusters. The Ct1 cluster during peak of fibrosis d14 displayed a decrease in the LIF signature compared to saline. However, during resolution, at d30 and d60, the LIF signature is enriched in the Ct1 cluster. Similarly, the Ct2 cluster in saline displays a high LIF signature. Upon bleo injury the signature decreases at d14 and recovers during resolution to its saline level.

Correlation analysis comparing *Cthrc1+* Ct1 to Ct2 at d14 and at d30 also confirms that the expression of canonical markers of LIFs such as *Limch1*, *Inmt*, *Macf1*, *Apoe* and *Tcf21* are upregulated at d30 (Figure 4c), therefore supporting a MYF to LIF transition. IPA® canonical pathways enrichment analysis of Ct2 at d14 and d30 compared to saline d14 alveolar fibroblasts, showed the enrichment of pathways like PDGF, GP6, TGFβ and IL-17 at day 14, and their subsequent downregulation at d30 (Figure 4d).

### Comparison of young vs old alveolar fibroblast subpopulations during fibrosis formation and resolution

Next, we carried out a sub clustering of the alveolar fibroblasts. Like in our previous study, alveolar fibroblasts can be subdivided into Al1, Al2 and Al3 (Figure 5a). Figure 5b shows the relative frequencies of the respective sub clusters, Al2 is the abundant cluster of the alveolar fibroblasts. Interestingly, at the peak of fibrosis (day 14), Al1 and Al2 show a decrease in their proportions which was partially, rescued during resolution at day 30 and 60. In contrast, Al3 proportion increased at day 14 and conversely decreased during resolution. These relative frequency changes for these clusters were not significantly observed in young mice (Figure S3c and d). We also did not observe a higher proportion of Al3 in old mice compared to young in saline d14 samples (Figure S3c).

**Figure 5:**
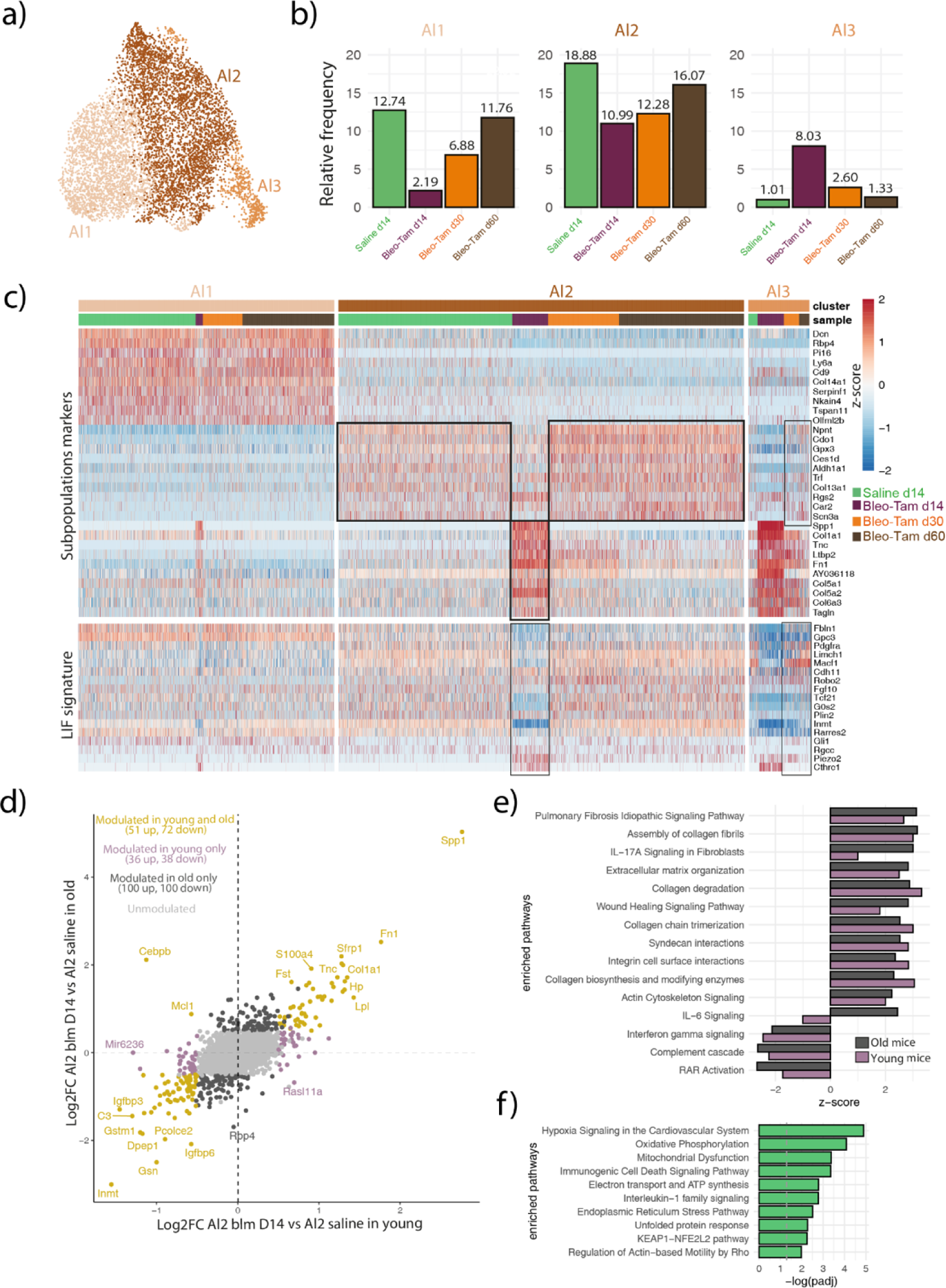
Comparison of alveolar fibroblasts subpopulations during fibrosis formation and resolution in old mice. a) UMAP of *Acta2+* cells isolated from saline d14 and Bleo-Tam d14, d30 and d60 lungs contributing to 3 alveolar fibroblasts subpopulations (Al1, Al2, Al3). b) Relative frequencies of each sub-populations at each time-point. **c)** Heatmap showing the scaled expression of alveolar fibroblasts subpopulations markers (top) and genes of the lipofibroblast (LIF) signature (bottom) in saline d14 and Bleo-Tam d14, d30 and d60 lungs. **d)** Correlation of the two log2 Fold change obtained by comparing Al2 of Bleo-Tam d14 sample to Al2 of saline d14 sample from old (52-56 weeks) and young (8-12 weeks) mice. Golden dots correspond to genes significantly modulated (|log2FC|>0.5 and padj < 0.05) in both old and young. Dark grey and mauve dots correspond respectively to genes only significantly modulated in old or in young. **e)** IPA® Canonical Pathways enrichment obtained from the differential analyses of Al2 of Bleo-Tam d14 sample compared to Al2 of saline d14 sample in both old and young. **f)** IPA® Canonical Pathways enrichment obtained from the upregulated genes in Al2 of saline d14 sample from old mice compared to Al2 of saline sample from young mice.

Figure 5c shows the heatmap of the most expressed gene markers for each Alveolar fibroblast sub cluster. At d14, following, bleo injury and at the peak of fibrosis, the Al2 cluster shows a decrease in its native signature and acquires the expression of fibrotic gene (*Spp1*, *Col1a1*, *Tnc, Ltbp2* and *Tagln*) adopting Al3 characteristics. In contrast, during resolution, Al2 cells retrieve back to native/saline state by losing the fibrotic gene expression. At the peak of fibrosis day 14, the LIF signature in Al2 cells show a decrease which is recovered during fibrosis resolution. Al1 cells also show a similar trend (acquisition of Al3 characteristics and decreased LIF signature following bleo at d14).

The Al3 cluster, which is quantitatively enriched following bleo at d14,displays a high levels of fibrotic genes expression (*Spp1*, *Col1a1*, *Tnc, Ltbp2* and *Tagln*). In young mice, our initial analysis indicated that Al3 cells, are mostly observed in Bleo-Tam d14 but not in Tam-Bleo d14. This indicates that these cells were initially *Acta2*-. However, during fibrosis formation, they acquire *Acta2* expression and are captured by our lineage tracing approach as *Acta2*+ cells. We proposed that these cells differentiate towards *Cthrc1*+Ct1 cells (Lingampally et al., 2024 in revision at ERJ). During resolution, the Al3 cluster shows a decrease in its fibrotic signature and shifts towards an Al2 phenotype. Similarly, the decreased LIF signature in this cluster at day 14 is also rescued during fibrosis resolution day 30 and 60 (Figure 5c).

It is therefore likely that Al2 *Acta2*- cells are a primary source for the Al3 *Acta2*+ cluster following bleo injury. Correlation analysis comparing Al2 clusters of Tam-Bleo d14 vs saline d14 in young and old mice, shows the activation of Al2 cluster with fibrotic genes (*Spp1*, *Sfrp1, Col1a1*, *Tnc* and *Fn1*) (Figure 5d).

At the transcriptomic level and in our experimental conditions, direct comparison between old and young samples was difficult to perform because of the differences in the UMI content between the two sets of experiments (Figure S3e). Nevertheless, in fibrotic conditions, an indirect comparison can be performed by confronting the log_2_FC in Al2 between Bleo-d14 vs saline d14 in old and young mice (Figure 5d). Overall, IPA® canonical pathways enrichment analysis indicated some age-dependent modulations (Figure 5e). Interestingly, we found that IL-17A and IL-6 signaling were both upregulated in Al2 old vs young (Figure 5e). To check for age-related differences in saline d14, we focused on upregulated genes in old compared to young Al2 cells. The genes upregulated in young versus old Al2 cells were not analyzed due to the lower sequencing depth in old mice resulting in a limited number of genes detected. Our analysis indicated the enrichment of several pathways in Al2 cells, such as hypoxia signaling, oxidative phosphorylation, IL-1 signaling, regulation of Actin-based motility by Rho, which could be relevant for the defective resolution in old mice (Figure 5f).

### Characterization of *CTHRC1*+ population in the human IPF lungs

To conduct an in-depth analysis of *CTHRC1*+ cells in human lungs, we combined the stromal compartment of healthy and IPF donors from various published datasets (described in the material and methods section). The resulting list of samples included 62284 cells in total (26391 control and 35893 IPF cells).

Following integration and dimension reduction, we identified 10 clusters (Figure 6a) including *LIMCH1+* lipofibroblasts (*LIMCH1*+LIF), *PLIN2*+ lipofibroblasts (*PLIN2*+LIF), activated aMYFs, *PI16+* adventitial fibroblasts, *DIO2+* peribronchial fibroblasts, MYFs, fibromyocytes (FibroM), *MYH11+* VSMC, *RGS5+* pericytes and *ITLN1+* mesothelial cells. Figure 6b shows the Dot plot for the marker genes for each subpopulation. UMAP plots showing the expression of marker genes for each subpopulation (Figure S4) and the LIF markers (Figure S5) in control and IPF lungs are also provided.

**Figure 6:**
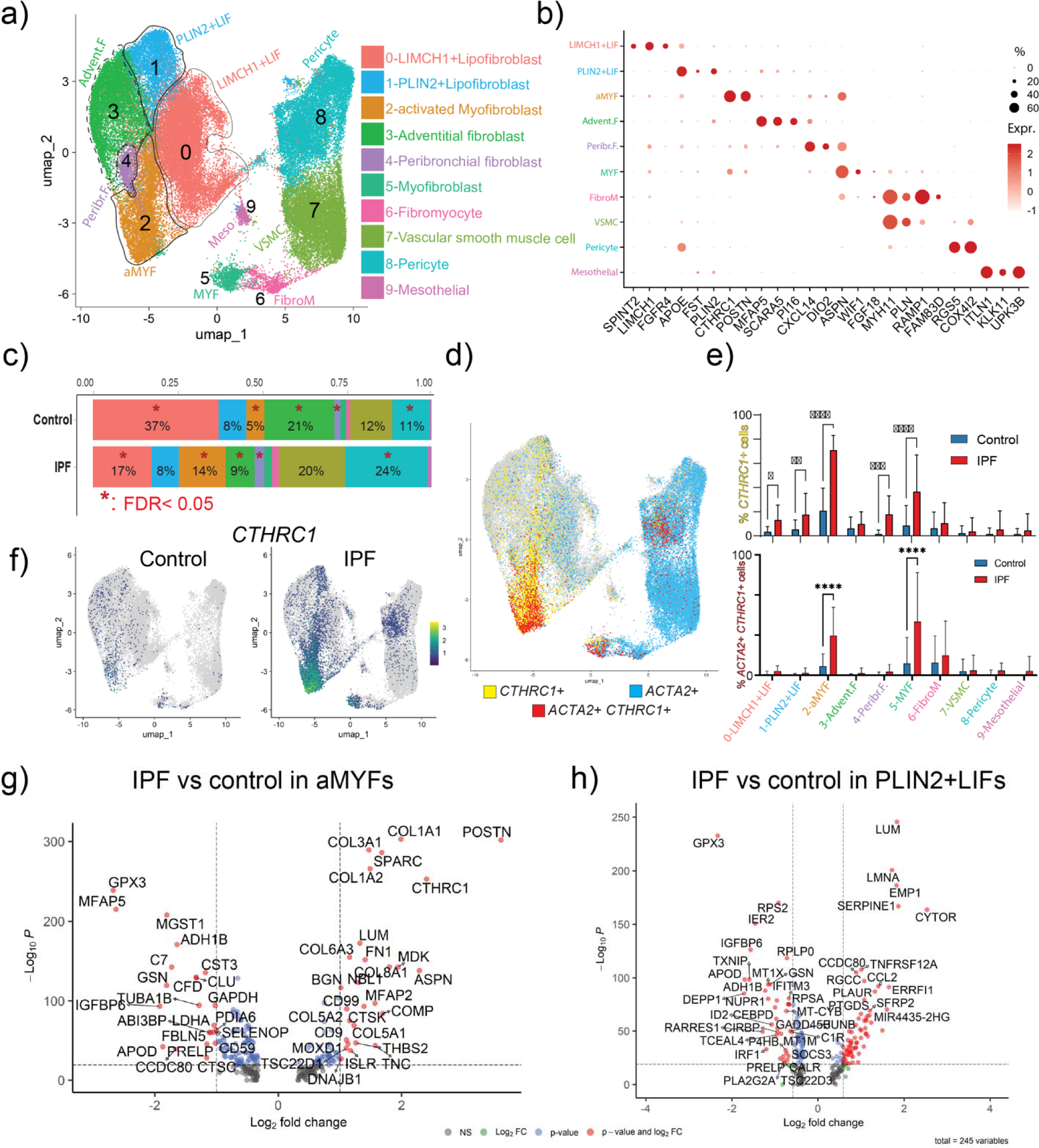
Characterization of the *CTHRC1*+ myofibroblasts in human lung stromal populations. a) UMAP plot for the integrated scRNAseq data of human lung stromal cells. b) Dot plot for the expression of established marker genes for stromal subpopulations of the human lung. Color gradient correlates with the average expression level while the size of dots indicates the percentage of cells expressing the corresponding gene. c) Bar plots indicating cell proportions in control and IPF cell clusters, respectively. Clusters significantly (FDR <0.05) affected by the disease are indicated with an asterisk. d) UMAP plot showing cells positive for *CTHRC1* (yellow) or *ACTA2* (blue) and positive for both (red) e) Bar-plot showing proportion of cells positive for *CTHRC1* and cells double positive for *CTHRC1* and *ACTA2* in each cluster. f) UMAP plot for the expression of *CTHRC1* in human control and IPF cells. The gradient color correlates with *CTHRC1* expression level (blue is low; yellow is high). g-h) Volcano plot of the genes differentially expressed between IPF and control in aMYFs (**g**) and PLIN2+LIFs (**h**). Grey dots correspond to genes that did not pass our fold-change (FC =2) and adjusted p-value (p-adj) cut-offs (10e-20). Genes with insufficient FC are shown in blue. Red dots are labeled with names for genes that met all requirements ( FC> 2 or FC <0.5 and padj < 10e-20).

The relative proportions of the different clusters are shown in Figure 6c. Interestingly, the proportion of *LIMCH1+*LIF and adventitial fibroblasts decreases significantly in IPF lungs compared to control donors, but not *PLIN2*+LIF. The transcriptional profile of *PLIN2*+LIF was highly altered in IPF lungs, with the upregulation of *LUM, EMP1, CYTOR* and the downregulation of *GPX3* for instance (Figure 6h). Fibromyocytes shared *MYH11/PLN* expression with VSMC and, similar to myofibroblasts, expressed *ASPN* (Figure 6b). This cluster also included a *FAM83D*+SMC subset that was first described in the Human Lung Cell Atlas (HLCA) study (Sikkema et al., 2023). MYFs were identified using the canonical markers *ASPN* and *WIF1*, while aMYF broadly expressed high levels of *CTHRC1* and *POSTN* (Figure 6b). Strikingly, the proportion of MYF and FibroM was comparable between control and IPF, but the population of aMYF was almost tripled in IPF lungs (Figure 6c), thus confirming that specific aMYF expansion is the major pathological change affecting non-vascular muscle cells in human lungs. *ACTA2+* cells are enriched in VSMCs and pericytes clusters (Figure 6d). Similar to our findings in mouse lungs ectopic low expression of *CTHRC1* was detected in clusters other than aMYFs, particularly pericytes and LIFs (Figure 6d). Additionally, a massive increase of *CTHRC1+* cells in IPF lungs was observed (Figure 6d-f). Interestingly, *ACTA2+ CTHRC1+* double positive cells are increased in IPF lungs and these cells are majorly emerging from aMYFs (Figure 6d, e). As expected, compared to healthy controls, *CTHRC1* was significantly upregulated in IPF-aMYF, along with ECM components (*POSTN*, collagens, *FN1*, *LUM*) and *SPARC* that promotes TGF-β-induced fibrosis (Carvalheiro et al., 2020) (Figure 6g).

Signature scores of Alveolar and cthrc1 clusters from old (Figure S6) and young mice (Figure S7) in human UMAPs were also carried out. Our results using the signatures identified in old mice (this study) indicated that the Al1 signature is enriched in Adventitial Fib. and *PLIN2*+LIF (Figure S6c) while the Al2 signature is enriched in *LIMCH1*+LIF (Figure S6d). Interestingly, the Al3 signature in mostly found in aMYFs supporting the proposed LIF to aMYF differentiation (Figure S6e). Ct1, Ct3 and Ct4 signatures were found enriched in *LIMCH1*+LIF and aMYF while the Ct2 signature was almost exclusively found in aMYF (Figure S6g-j). Finally, the LIF markers APOE, FST and PLIN2 were enriched in the *PLIN2*+LIF while the overall LIF signature used in mice was mostly enriched in the *LIMCH1*+LIF (Figure S6b, f). Similar results were obtained using the signatures identified in young mice (Figure S7).

### Transcriptional programs associated with lipofibroblast differentiation into *CTHRC1*+ myofibroblasts in human IPF

Next, to predict the possible origin of aMYFs in human lungs, we performed a trajectory analysis on the stromal compartment using Slingshot (Street et al., 2018). The resulting lineage ordered the fibroblast clusters as follows: adventitial FIB > *PLIN2+*LIF > *LIMCH1*+LIF > aMYF (Figure S8a). To gain more insights into the mechanisms involved in the inferred LIF-to-aMYF transition, we then focused our trajectory analysis on the cells belonging to the *PLIN2+*LIF, *LIMCH1*+LIF and aMYF clusters. Cell’s transcriptional progress along the LIF-to-aMYF trajectory was evaluated by a pseudotime (Figure 7a, S8b) and split into 3 major stages (Figure 7b). *PLIN2*+LIFs (stage 1) and aMYFs (stage 3) from IPF patients were spread onto a larger range of pseudotimes, suggesting a higher heterogeneity of cell states in these clusters (Figure 7b). We found 1164 and 1385 genes significantly associated with cell pseudotime in control and IPF lungs, respectively, of which 486 genes were common to both conditions (Figure 7c). In addition, many of the genes associated with pseudotime in control cells belonged to the same enriched ontology term as genes specific to IPF cells (Figure 7c). Comparison of the smoothed expression of all 2063 genes associated with pseudotime in either control or IPF cells showed that, compared to control cells, cells of IPF lungs cells tend to upregulate large sets of genes, especially at early pseudotimes of stage 1 (*PLIN2*+LIF) and later pseudotimes of stage 3 (aMYFs) (Figure S8c). Together this suggests that control and IPF cells share similar gradual transcriptional changes during the LIF-to- aMYF transition; however, IPF cells exhibit a hyperactivation of some of these biological pathways.

**Figure 7:**
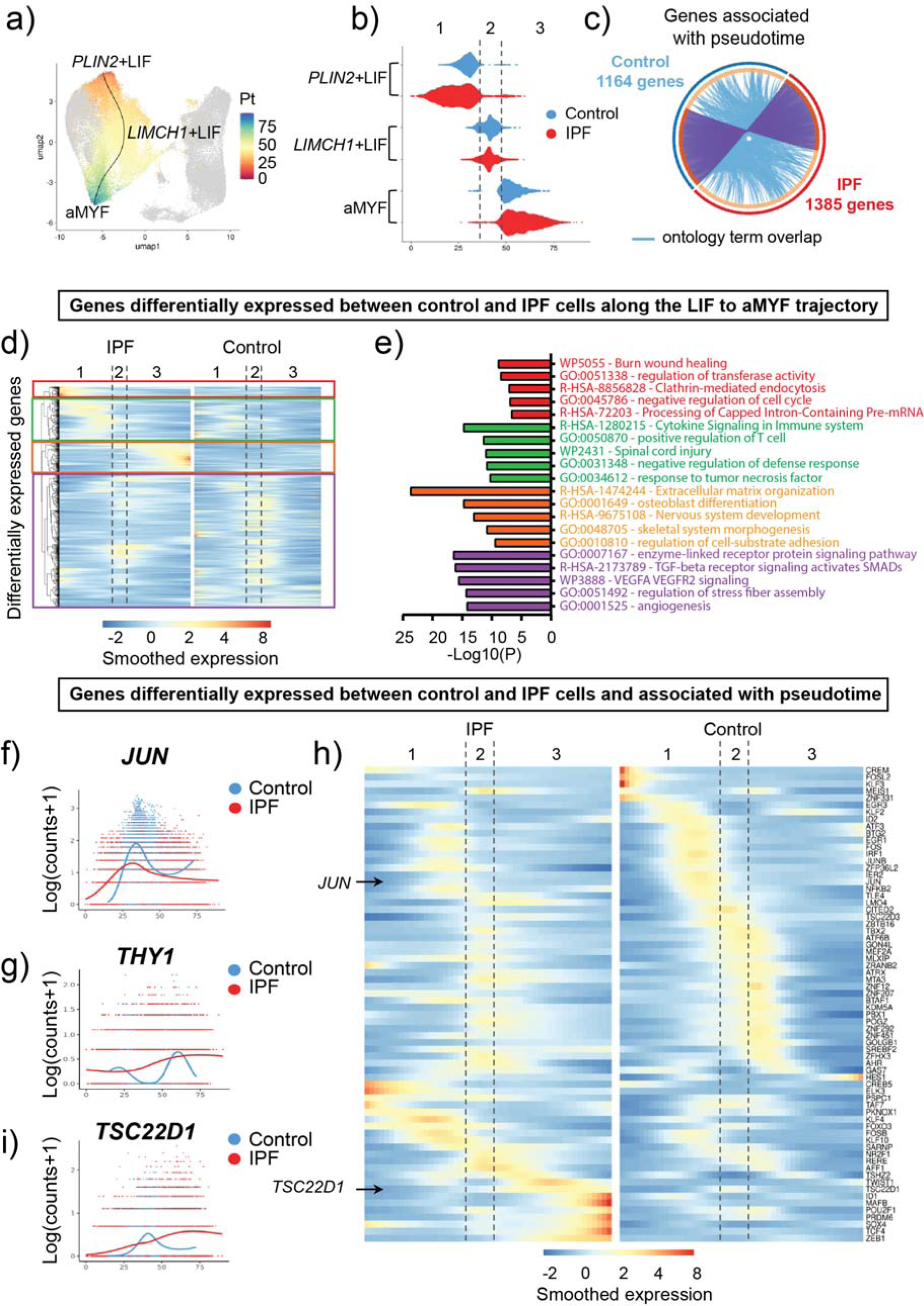
Inferred trajectory for the differentiation of human lipofibroblast into *Cthrc1*+ myofibroblasts. a) The trajectory inferred on the *PLIN2*+ LIF- *LIMCH1*+ LIF- aMYF subset was projected onto the UMAP dimension reduction plot (black line). The gradient color indicates cell pseudotime (red is low, blue is high). **b)** Ordering of cells by pseudotime Dotted lines delineate the 3 major stages. **c)** Circos plot for the lists of genes associated with pseudotime in control (1164 genes) and IPF cells (1385 genes). The 486 genes shared by both conditions are shown by dark orange arcs and linked by purple lines. Light orange arcs represent specific genes. Blue lines link the genes, that fall under the same ontology term (statistically significantly enriched and with size no larger than 100). **d)** The joint clustered heatmap of the smoothed expression of the 766 genes differentially expressed along the LIF-to-aMYF trajectory between control and IPF cells revealed 4 main clusters of expression patterns. Expression values are ordered according to pseudotime from left to right and cell stages are marked by dotted lines. **e**) Genes of each pattern were submitted to pathway analysis using Metascape. The top-5 of enriched biological pathways are shown and colored by the cluster of genes they correspond to (d). **f**-**g,i**) Scatter plots for the gene expression measures against cell pseudotime and their respective smoothed expression values in each condition (control and IPF are shown in blue and red, respectively). **h**) Heatmap of the smoothed expression for transcription factors significantly associated with pseudotime in the LIF-to-aMYF trajectory, differentially expressed between the start and the end of the trajectory, and also between control and IPF cells.

To gain a better insight into the differences between patient’s cells along the LIF-to- aMYF trajectory, we studied the genes whose expression patterns were significantly different in control and IPF cells. The resulting 766 differentially expressed genes were clustered into 4 main groups according to their expression pattern along the pseudotime (Figure 7d). Pathway enrichment analysis (Figure 7e) showed that later stages of IPF *PLIN2*+LIFs upregulated gene sets related to inflammation and immunity (green). Genes activated at the *LIMCH1*+LIF to aMYFs transition (violet) seemed for the most part to be more actively transcribed in normal cells compared to IPF cells. This large cluster was enriched in genes were involved in signal transduction including TGF-β- signaling (Figure 7e), with the transcription factor *JUN* being the most significant in this cluster of genes (Figure 7f, S8d). This suggests that pro-fibrotic processes may be activated in homeostatic conditions in *LIMCH1*+LIFs and that IPF-related abnormalities are mainly found at later pseudotimes (stage 3). As expected, the end of the trajectory was characterized by the upregulation of *CTHRC1* (Figure S8e) and genes related to ECM remodeling and muscle differentiation in IPF aMYFs (orange, Figure 7e). The most significant gene of this group was *THY1* whose expression level was sustained in IPF aMYFs, as compared with the transient activations during state transitions in control cells (Figure 7g). IPF upregulated *THY1* in several fibroblast subsets (Figure S8d).

To identify the drivers for the transcriptional programs associated with the LIF-to-aMYF transition and aberrantly activated in IPF lungs, we analyzed the expression patterns of the transcription factors that were significantly associated with pseudotime and also differentially expressed between control and IPF cells (Figure 7h). Among them, a few transcription factors were upregulated in IPF cells, mostly at stage 3 corresponding to aMYFs (Figure 7i). We thus found the gene *TSC22D1*, which showed an increasing and sustained transcription over the trajectory in IPF cells, thus contrasting with the transient activation observed in control *LIMCH1+*LIFs (Figure 7h, S8d). *TSC22D1* has been described as a potentiator of TGF-β-signaling and thus, contributes to myocardial fibrosis (Yan et al., 2011). Therefore, the aberrant expression of *TSC22D1* may promote aMYF terminal differentiation and the resulting fibrosis.

## Discussion

IPF is a lethal lung disease caused by irreversible fibrosis and associated with aging. Considering these later criteria, it has been proposed that old mice rather than young mice represent a much more relevant model to study IPF due to elevated fibrosis and delayed resolution following bleo exposure (Caporarello et al., 2020; Jenkins et al., 2017; Redente et al., 2011).

The underlying causes for such difference in the response to bleomycin-induced fibrosis in old versus young mice are progressively emerging. Among them, others and we have proposed that the nature of the lung mesenchyme is changing. Our study provides evidence for a LIF to MYF-like differentiation process during aging. The alveolosphere assay has been instrumental to evaluate the activity of the resident mesenchymal cells in supporting the proliferation of AT2 stem cells. Using this model, we show that this niche activity is lost already in 8 months-old animals, serving as a rational to use 12 months-old animals for our studies instead of the 1.5-2 years old mice that are currently used for studies involving aging. In 12 months-old mice, we observed increased fibrosis formation and delayed fibrosis resolution compared to 2 months-old animals for the same dose of bleo.

Interestingly, we previously reported that the rMC Sca1+ niche activity was also impacted by obesity and by gender, with *ob/ob* versus wild type (WT) mice and male versus female WT mice displaying the lower niche activity (Taghizadeh et al., 2021). These observations made in mice are consistent with what is observed in humans where COVID-19-infected aged males suffering from metabolic diseases are more prone to develop pneumonia (Cai et al., 2020; Muniyappa & Gubbi, 2020; Nouri- Keshtkar et al., 2021; Zhang et al., 2020).

Recently, the analysis of the pathways at the transcriptomic and proteomic level associated with fibrosis appear to be similarly regulated between young and aged mice (Klee et al., 2023). A similar observation was done following the analysis of metabolomic and lipidomic changes in old versus young animals following bleo injury (Weckerle et al., 2022).

However, in old mice an increase inflammatory state was detected (Klee et al., 2023). Interestingly, we noticed a persistence of IL-1, IL-17A and IL-6 signaling in old Al2 cells in saline and after bleo injury (please see model in figure 8) supporting a dysfunctional regulation of the immune response in old animals. Further studies will have to be done to understand the functional of the immune response in the context of aging and fibrosis formation.

**Figure 8:**
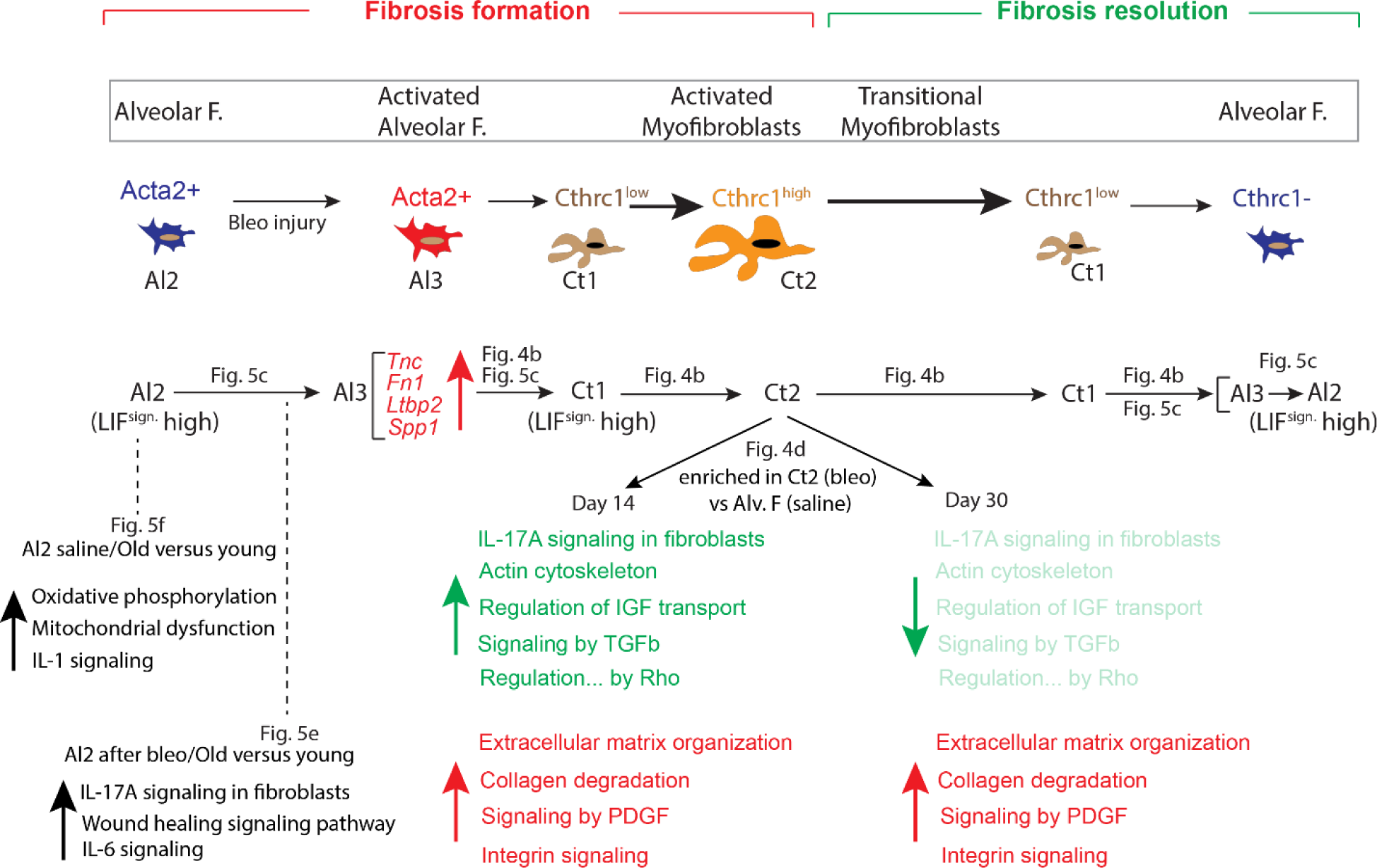
Model of LIF to MYF reversible switch during fibrosis formation and resolution in old mice. The main contributors during fibrosis formation to *Cthrc1*+ aMYF (Ct1-Ct4) are Acta2+ LIF^high^ Alveolar Fibroblasts (Al2) which get activated and differentiate to Al3 to express a fibrotic signature (*Spp1*, *Tnc*, *Ltbp2* and *Fn1*). These activated Al3 cells eventually differentiate into *Cthrc1*^Low^ LIF^high^ aMYF belonging to the Ct1 cluster. These Ct1 cells further differentiate massively to contribute to Ct2 *Cthrc1*^high^ aMYFs. During fibrosis resolution, the opposite occurs with Ct2 cells decreasing their signature and differentiating into Ct1 cells, which eventually differentiate back to the Al3 cluster and finally to *Cthrc1*- LIF^high^ Al2 Alveolar fibroblasts cluster. The pathways modulated in Al2 saline (old versus young), Al2 after bleo (old versus young), and enriched in Ct2 (bleo) vs Alv. F (saline) are shown.

Overall, our lineage tracing if the aMYF indicates that the basic cellular mechanisms are conserved between old and young mice. Our results indicate an alveolar fibroblast to a *Cthrc1*+aMYF reversible switch during fibrosis formation and resolution (see model in Figure 8). Interestingly, we identified pathways which are downregulated in *Cthrc1*+aMYF (cluster Ct2) between Bleo-Tam d30 and Bleo-Tam d14. These pathways include IL-17A and signaling by TGFb which are major contributors to fibrosis formation. Such downregulation over time indicates that the resolution process is taking place in old mice. However, we found also other signaling pathways such as signaling by PDGF, integrin signaling and collagen degradation which are still highly active at day 30, consistent with the observed persistence of fibrosis.

Interestingly, we have previously reported that metformin, an anti-diabetic drug also thought to have an impact on longevity (Barzilai, Crandall, Kritchevsky, & Espeland, 2016), is capable of triggering MYF to LIF differentiation in vitro (Kheirollahi et al., 2019) (Pacheco et al., 2023) and improve fibrosis resolution in vivo in mice (Kheirollahi et al., 2019; Rangarajan et al., 2018). The clinical use of metformin in combination with the gold standard treatments (pirfenidone or nintedanib) to attenuate fibrosis is still at its enfancy but highlights how enhancing the MYF to LIF differentiation process could be used clinically to provide new therapeutic treatments for IPF patients. In the future, new drugs capable of triggering LIF differentiation, with or without an impact on the fibrotic profile are likely to be instrumental in restoring normal lung function and could be identified using our recently described in vitro model using WI-38 human embryonic lung fibroblasts (Pacheco et al., 2023).

Finally, our extended analysis of the status of the mesenchymal cells in human lungs including all the available scRNA-seq datasets supports the LIF to aMYF differentiation and validates the enrichment of the *Cthrc1*+ Ct2 signature initially found in mice in the aMYFs cluster.

In conclusion, our data obtained in old mice, support the results obtained in young mice concerning the LIF-MYF reversible switch. We propose that functionally targeting this switch by increasing LIF differentiation and/or decreasing MYF differentiation will be a promising approach for innovative IPF treatments. The basic pathways activated in the Ct2 cluster during fibrosis formation and resolution identified in our studies in young and old mice will be a starting point to identify potential therapeutic drugs.

## Acknowledgements

We thank Kerstin Goth for the animal husbandry and genotyping. We also thank Dr. Thomas Sontag and Dr. Ingrid Henneke for their help with the animal protocols and ethical approvals.

## Author contributions

AL contributed to the conceptualizing of the project, writing the animal protocols, carrying out the experiments, analysis and interpretation of the data and writing the manuscript. MT carried out the bioinformatic analysis on the generated mouse samples. OM, VD carried out the data mining on the human IPF samples. SHa, helped with the bleo injury model. JK and AIVA performed the FACS and scRNA-seq analysis. SHe, HPM, CS, and EEA contributed to the conceptualization of the project. BM, CC, SB conceptualized the project, provided funding, analyzed and interpreted the data and wrote the manuscript. All authors read and agreed with the results presented in the manuscript.

---

**Figure S1:**
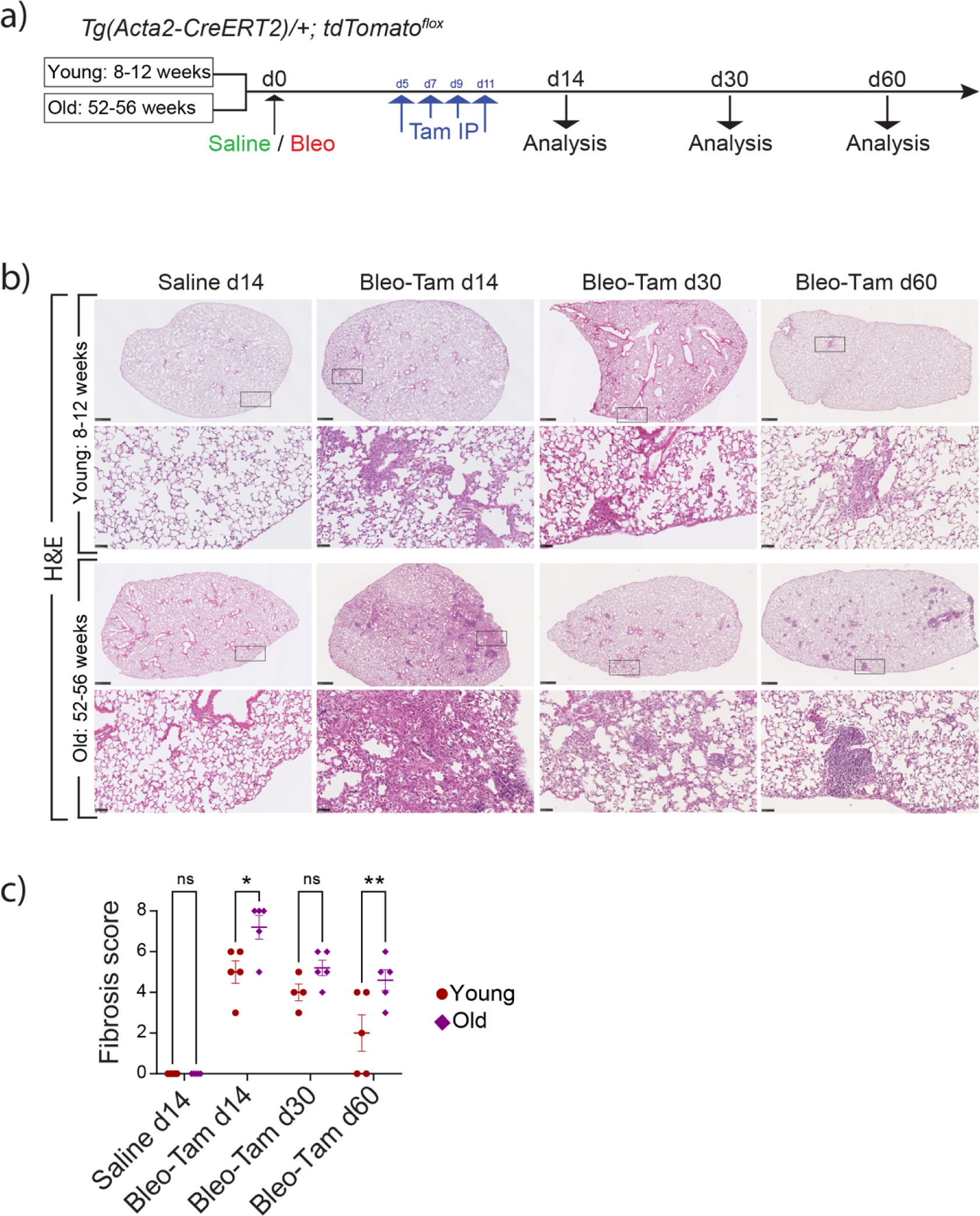
Comparison of fibrosis in young vs old mice. a) Female *Tg(Acta2- CreERT2)/+;tdTomato^flox^* Young (8-12 weeks) and old (52-56 weeks) mice are either subjected to saline or 2U/Kg Bleomycin (Bleo). Control lungs are collected at day14 following saline administration and experimental lungs (Bleo-Tam) are collected at d14, d30 and d60 following bleomycin injury. **b)** Corresponding low and high magnification of H&E staining showing massive fibrosis formation in old mice lungs compared to young at Bleo-Tam d14. Additionally, fibrotic lesions/regions are still present at resolution Bleo- Tam d30and Bleo-Tam d60 in formation in both young and old mice lungs. **c)** Comparison of the fibrotic scores between young and old mice at the different time points and conditions. Scale bar: b: Low magnification- 500µm, High magnification- 50µm; Statistical analysis was performed using: (c) 2-way ANOVA with Šídák’s post hoc test for multiple comparisons. *: P<0.05; **: P<0.01.

**Figure S2:**
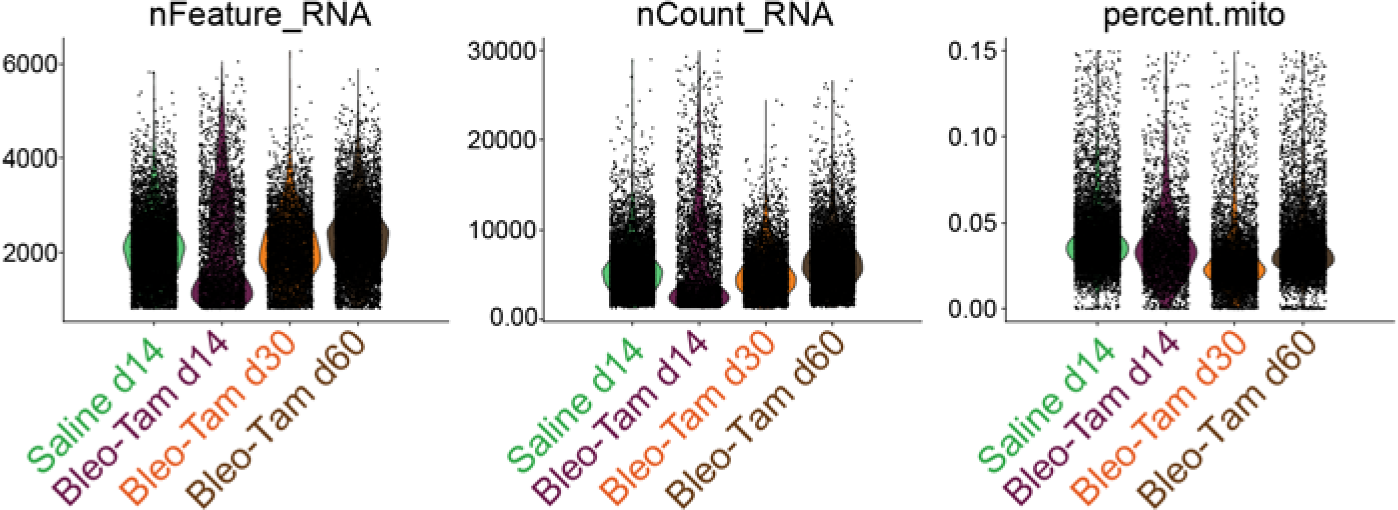
Quality control of scRNA-seq datasets from lineage-labeled *Acta2*+ (tdTom+) cells sorted from saline d14 and Bleo-Tam lungs at d14, d30 and d60 (52-56 weeks-old mice). nFeature_RNA (the number of genes detected in each cell), nCount_RNA (total number of RNA molecules detected within a cell) and the percentage of mirochondrial genes are quantified for each condition.

**Figure S3:**
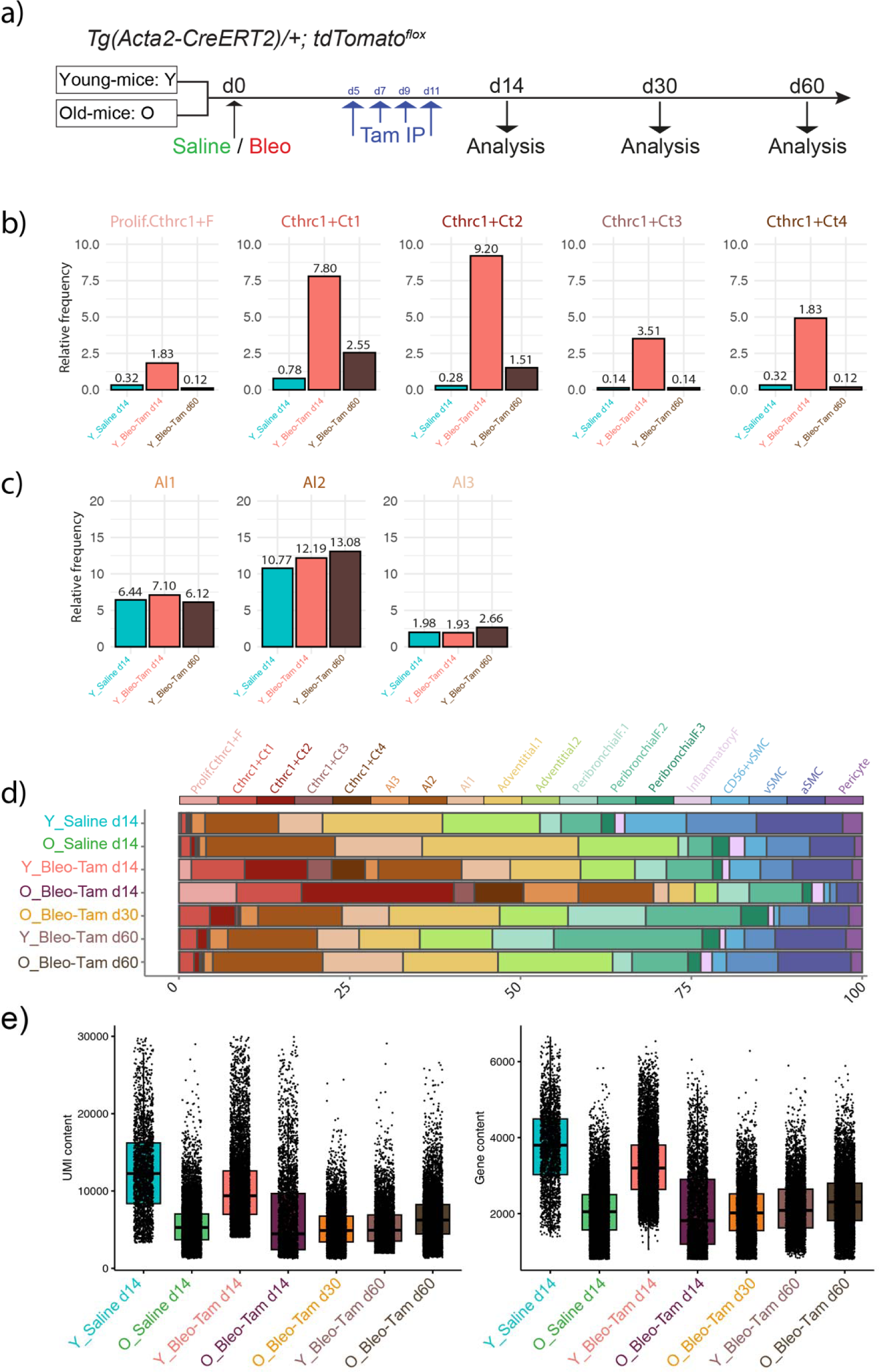
Comparison of Cthrc1 and Alveolar fibroblasts sub-clusters in young vs old mice a) Female *Tg(Acta2-CreERT2)/+;tdTomato^flox^* young (8-12 weeks) and old (52-56 weeks) mice are either subjected to saline or 2U/Kg Bleomycin (Bleo). Saline lungs are collected at d14 and experimental lungs (Bleo-Tam) are collected at d14, d30 and d60 following bleomycin injury. b) Relative frequencies of each *Cthrc1*-expressing subpopulations at each time-point in young mice. c) Relative frequencies of each alveolar fibroblasts subpopulations at each time-point in young mice. d) Relative frequencies of each subpopulation in each sample from the integrated dataset of young and old mice. e) UMI and gene content in each sample from the integrated dataset of young and old mice.

**Figure S4:**
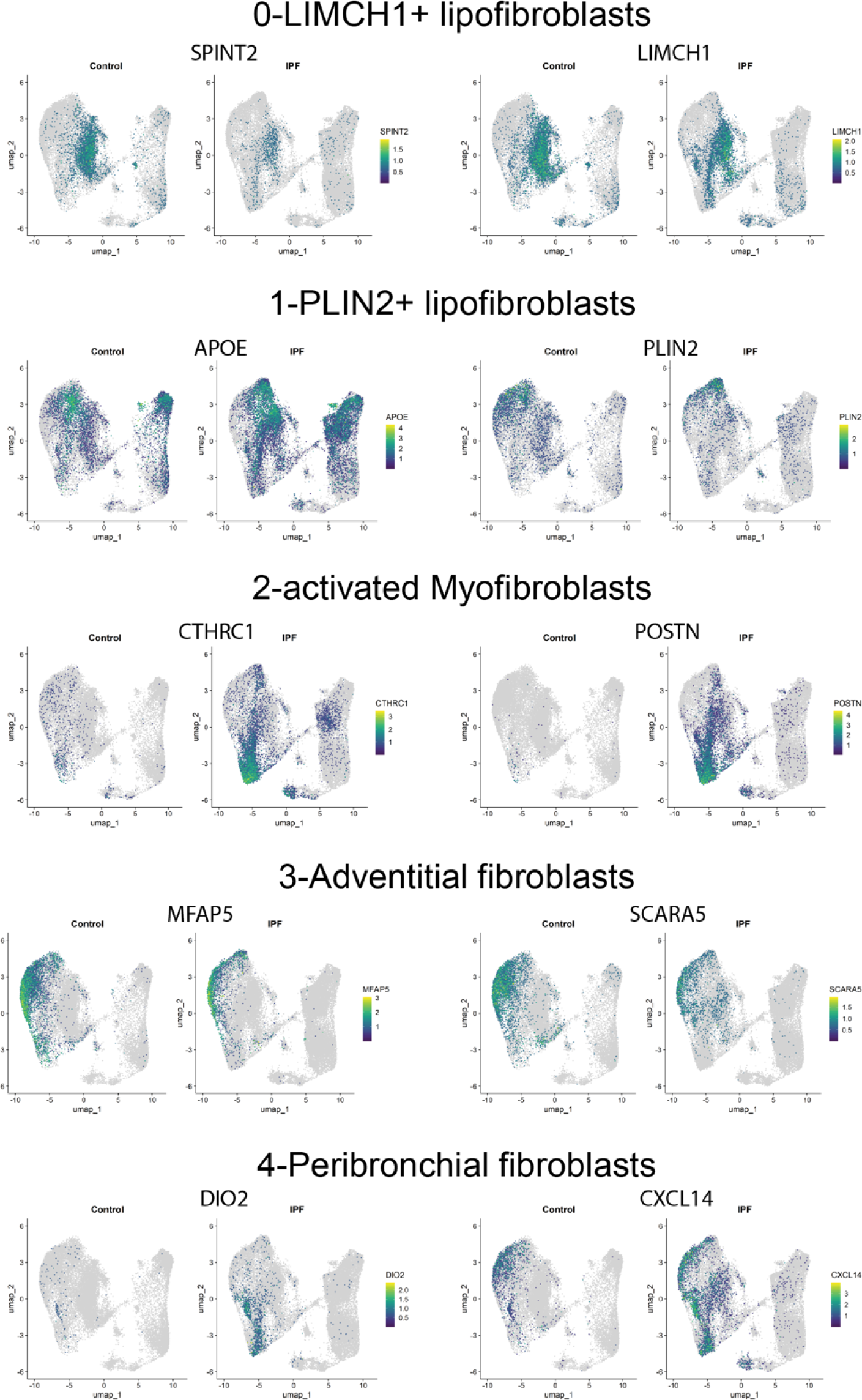
UMAP plot for the expression of marker genes for different clusters in human stromal cells between Control and IPF. The gradient color correlates with genes expression level (blue is low; yellow is high).

**Figure S5:**
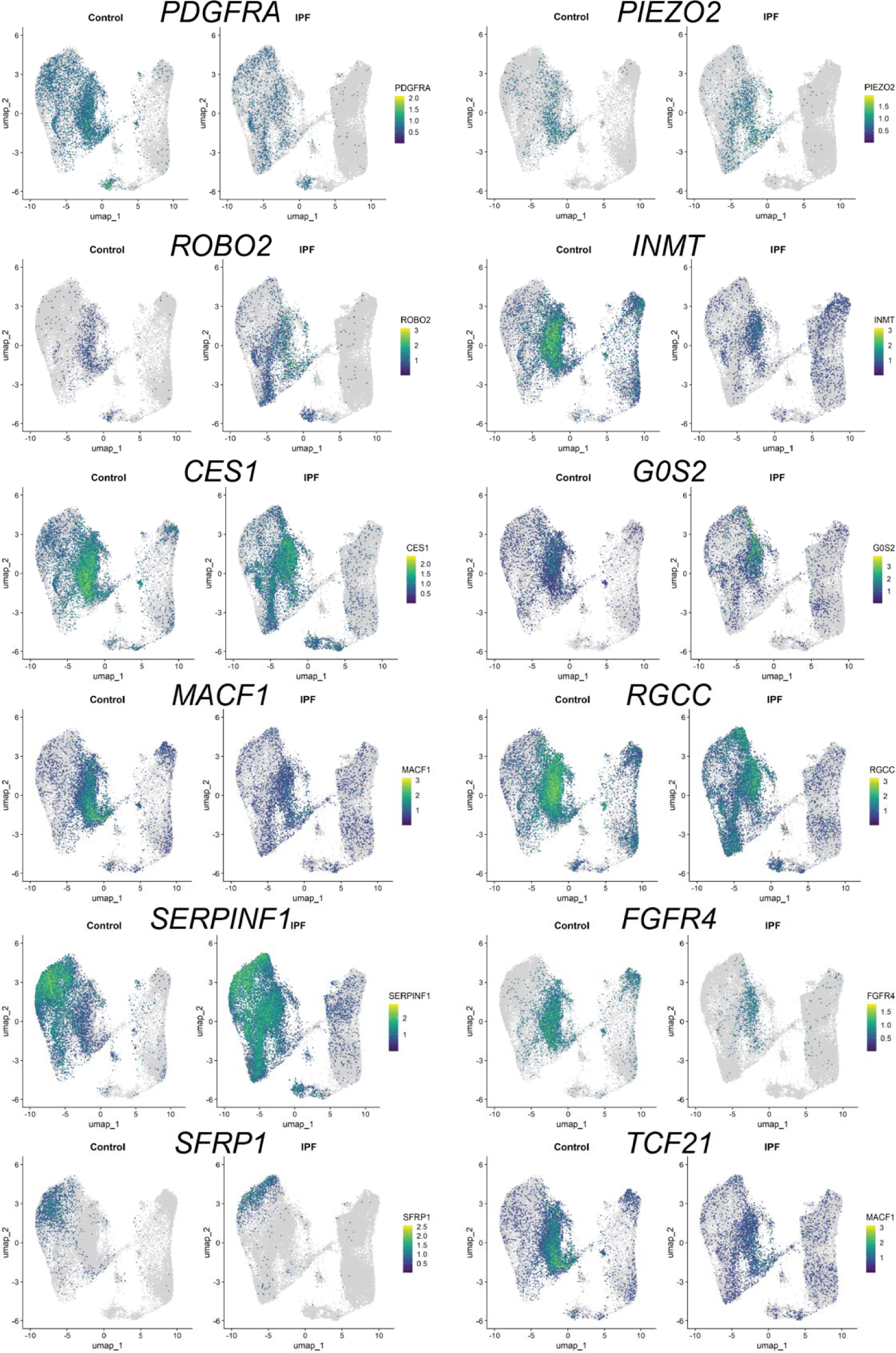
UMAP plot for the expression of LIF genes in human stromal cells between Control and IPF. The gradient color correlates with genes expression level (blue is low; yellow is high).

**Figure S6:**
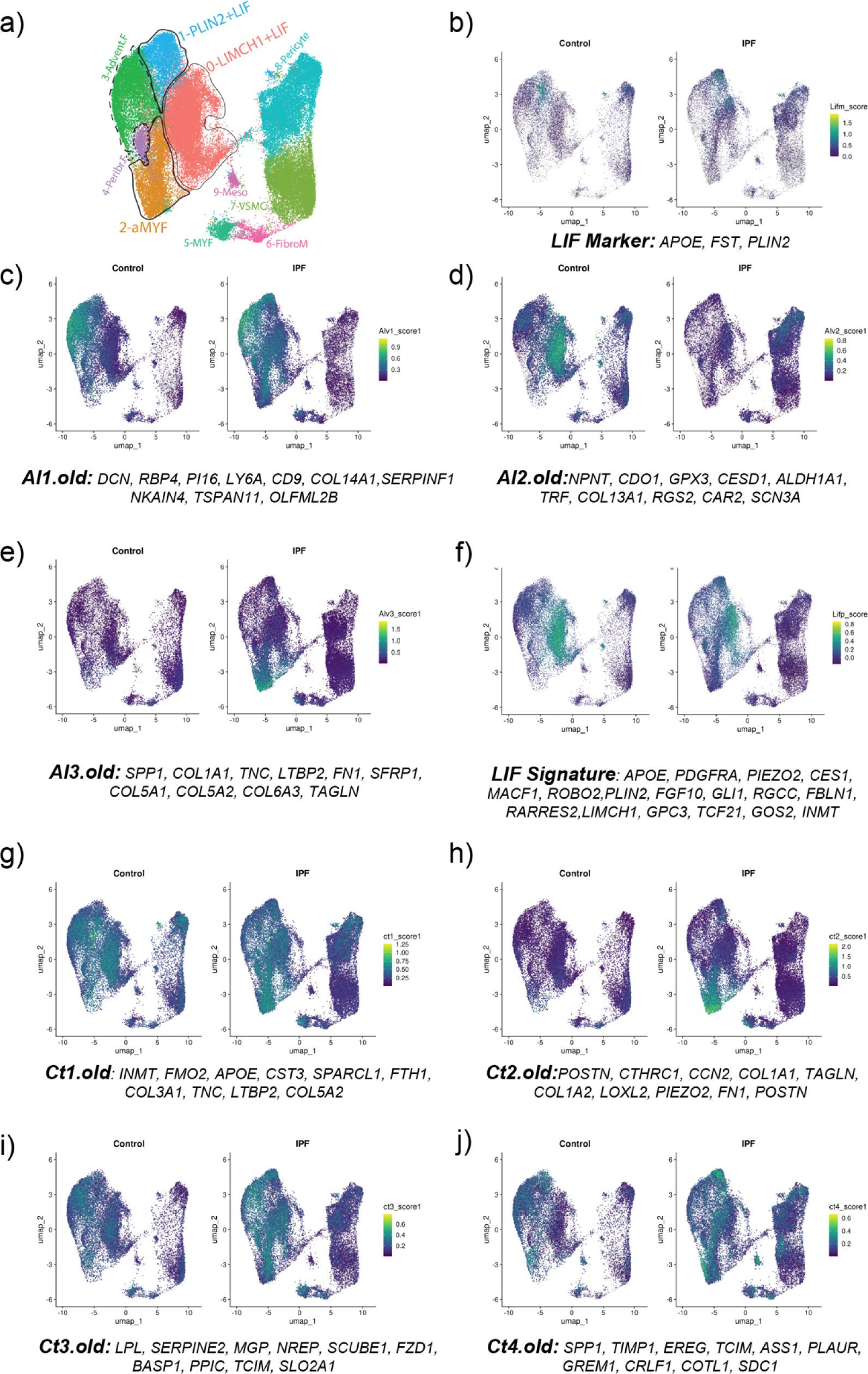
a) Integrated UMAP plot of human lung stromal cells. b-j) UMAP plot of the module score (expression of a gene program) for the following gene signature (identified in old Acta2mice): LIF markers (b), Al1 (c), Al2 (d), Al3 (e), LIF signature (f), Ct1 (g), Ct2 (h), Ct3 (i) and Ct4 (j). The list of genes included in each score is shown below each plot.

**Figure S7:**
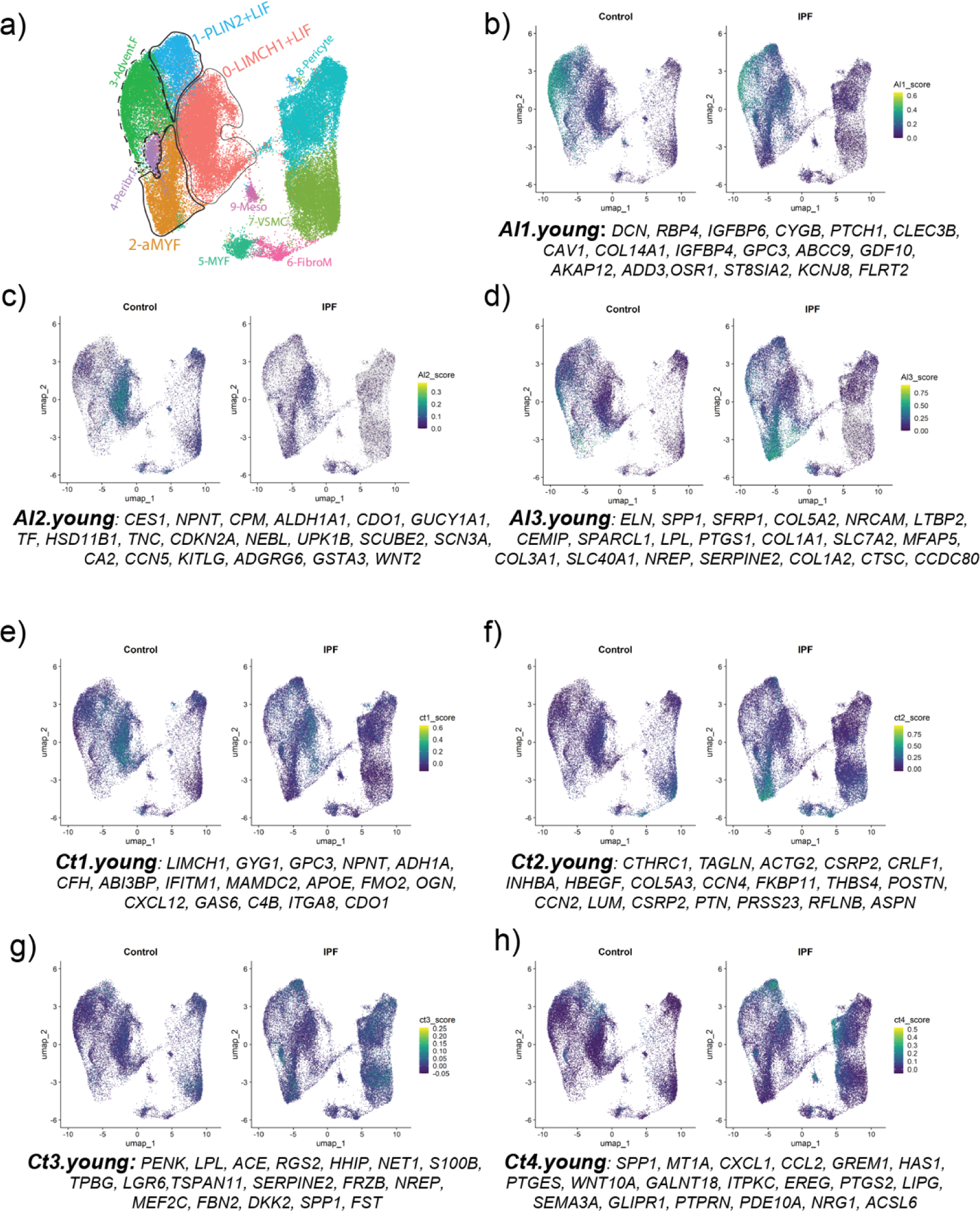
a) Integrated UMAP plot of human lung stromal cells. b-h) UMAP plot of the module score (expression of a gene program) for the following gene signature (identified in young Acta2mice): Al1 (b), Al2 (c), Al3 (d), Ct1 (e), Ct2 (f), Ct3 (g) and Ct4 (h). The list of genes included in each score is shown below each plot.

**Figure S8:**
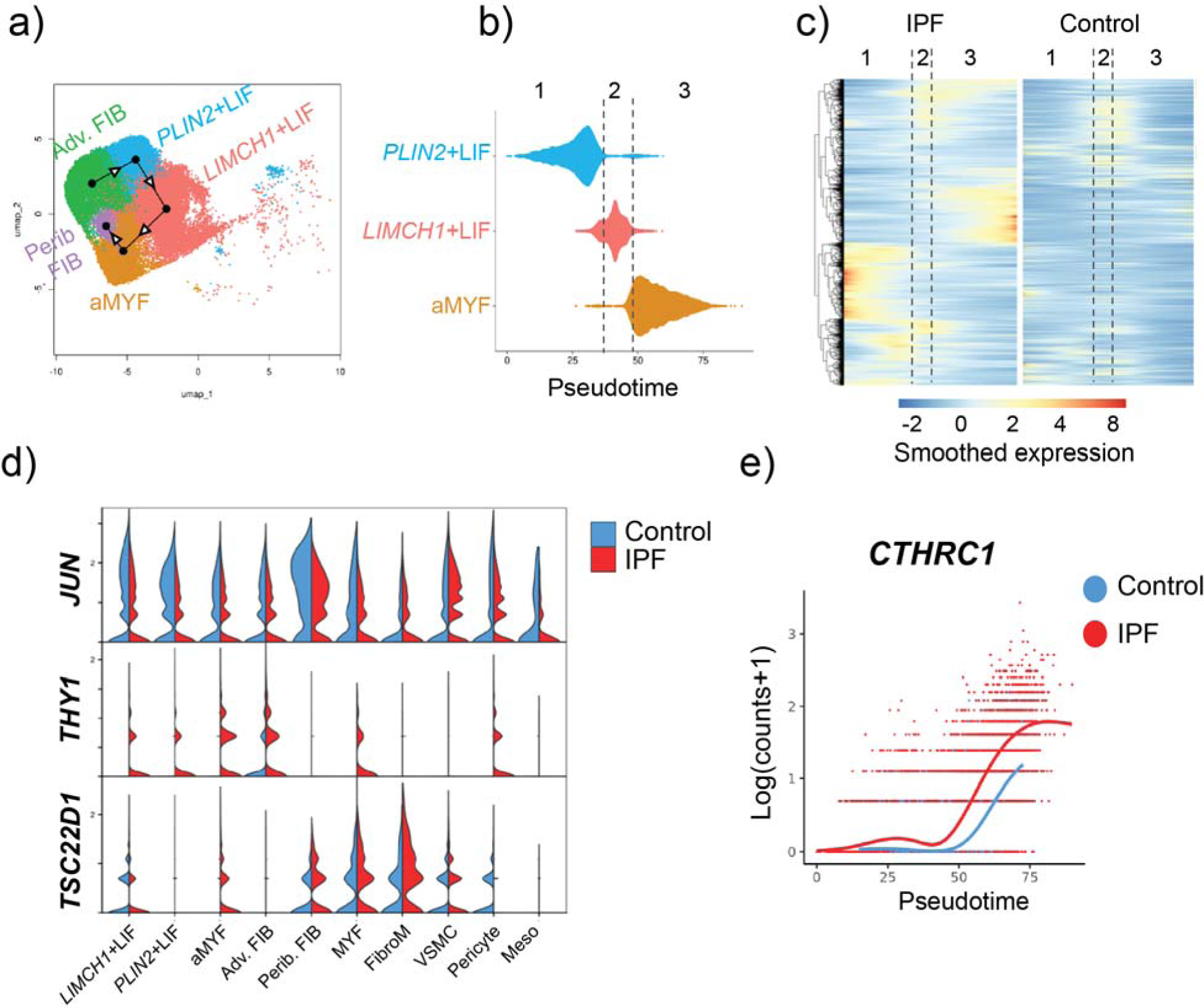
**a)** UMAP plot for the fibroblast subset submitted to Slingshot analysis and resulting trajectory originating from adventitial fibroblasts and ending in peribronchial fibroblast cluster. **b)** Cells of the *PLIN2*+ LIF, *LIMCH1*+ LIF, and aMYF were subsetted and ordered by pseudotime. **c)** Joint clustered heatmap of the smoothed expression of all 2063 genes significantly associated with pseudotime in either control or IPF samples in the LIF-to-aMYF trajectory. Expression values are ordered according to pseudotime from left to right. **d)** Violin plot for the normalized expression of *JUN, THY1* and *TSC22D1* in each cluster of control (blue) or IPF (red) human samples. **e)** Scatter plot for the expression measures of *CTHRC1* against cell pseudotime and its smoothed expression values in each condition (control and IPF are shown in blue and red, respectively).

## References

1. Adams, T. S., Schupp, J. C., Poli, S., Ayaub, E. A., Neumark, N., Ahangari, F., Kaminski, N. (2020). Single-cell RNA-seq reveals ectopic and aberrant lung-resident cell populations in idiopathic pulmonary fibrosis. Sci Adv, 6(28), eaba1983. doi:10.1126/sciadv.aba1983

2. Amara, N., Goven, D., Prost, F., Muloway, R., Crestani, B., & Boczkowski, J. (2010). NOX4/NADPH oxidase expression is increased in pulmonary fibroblasts from patients with idiopathic pulmonary fibrosis and mediates TGFbeta1-induced fibroblast differentiation into myofibroblasts. Thorax, 65(8), 733–738. doi:10.1136/thx.2009.113456

3. Ashcroft, T., Simpson, J. M., & Timbrell, V. (1988). Simple method of estimating severity of pulmonary fibrosis on a numerical scale. J Clin Pathol, 41(4), 467–470. doi:10.1136/jcp.41.4.467

4. Barzilai, N., Crandall, J. P., Kritchevsky, S. B., & Espeland, M. A. (2016). Metformin as a Tool to Target Aging. Cell Metab, 23(6), 1060–1065. doi:10.1016/j.cmet.2016.05.011

5. Bueno, M., Calyeca, J., Rojas, M., & Mora, A. L. (2020). Mitochondria dysfunction and metabolic reprogramming as drivers of idiopathic pulmonary fibrosis. Redox Biol, 33, 101509. doi:10.1016/j.redox.2020.101509

6. Cai, Q., Chen, F., Wang, T., Luo, F., Liu, X., Wu, Q., Xu, L. (2020). Obesity and COVID-19 Severity in a Designated Hospital in Shenzhen, China. Diabetes Care, 43(7), 1392–1398. doi:10.2337/dc20-0576

7. Caporarello, N., Meridew, J. A., Aravamudhan, A., Jones, D. L., Austin, S. A., Pham, T. X., Ligresti, G. (2020). Vascular dysfunction in aged mice contributes to persistent lung fibrosis. Aging Cell, 19(8), e13196. doi:10.1111/acel.13196

8. Carvalheiro, T., Malvar Fernandez, B., Ottria, A., Giovannone, B., Marut, W., Reedquist, K. A., Radstake, T. R. (2020). Extracellular SPARC cooperates with TGF-beta signalling to induce pro-fibrotic activation of systemic sclerosis patient dermal fibroblasts. Rheumatology (Oxford), 59(9), 2258–2263. doi:10.1093/rheumatology/kez583

9. Chanda, D., Rehan, M., Smith, S. R., Dsouza, K. G., Wang, Y., Bernard, K., Thannickal, V. J. (2021). Mesenchymal stromal cell aging impairs the self-organizing capacity of lung alveolar epithelial stem cells. Elife, 10. doi:10.7554/eLife.68049

10. Chung, M. P., Monick, M. M., Hamzeh, N. Y., Butler, N. S., Powers, L. S., & Hunninghake, G. W. (2003). Role of repeated lung injury and genetic background in bleomycin-induced fibrosis. Am J Respir Cell Mol Biol, 29(3 Pt 1), 375-380. doi:10.1165/rcmb.2003-0029OC

11. Cronkhite, J. T., Xing, C., Raghu, G., Chin, K. M., Torres, F., Rosenblatt, R. L., & Garcia, C. K. (2008). Telomere shortening in familial and sporadic pulmonary fibrosis. Am J Respir Crit Care Med, 178(7), 729–737. doi:10.1164/rccm.200804-550OC

12. Degryse, A. L., Tanjore, H., Xu, X. C., Polosukhin, V. V., Jones, B. R., McMahon, F. B., Lawson, W. E. (2010). Repetitive intratracheal bleomycin models several features of idiopathic pulmonary fibrosis. Am J Physiol Lung Cell Mol Physiol, 299(4), L442–452. doi:10.1152/ajplung.00026.2010

13. El Agha, E., Moiseenko, A., Kheirollahi, V., De Langhe, S., Crnkovic, S., Kwapiszewska, G., Bellusci, S. (2017). Two-Way Conversion between Lipogenic and Myogenic Fibroblastic Phenotypes Marks the Progression and Resolution of Lung Fibrosis. Cell Stem Cell, 20(2), 261–273 e263. doi:10.1016/j.stem.2016.10.004

14. Hao, Y., Hao, S., Andersen-Nissen, E., Mauck, W. M., 3rd, Zheng, S., Butler, A., Satija, R. (2021). Integrated analysis of multimodal single-cell data. Cell, 184(13), 3573-3587 e3529. doi:10.1016/j.cell.2021.04.048

15. Hao, Y., Stuart, T., Kowalski, M., Choudhary, S., Hoffman, P., Hartman, A., Satija, R. (2022). doi:10.1101/2022.02.24.481684

16. Jenkins, R. G., Moore, B. B., Chambers, R. C., Eickelberg, O., Konigshoff, M., Kolb, M., Molecular, B. (2017). An Official American Thoracic Society Workshop Report: Use of Animal Models for the Preclinical Assessment of Potential Therapies for Pulmonary Fibrosis. Am J Respir Cell Mol Biol, 56(5), 667–679. doi:10.1165/rcmb.2017-0096ST

17. Kheirollahi, V., Wasnick, R. M., Biasin, V., Vazquez-Armendariz, A. I., Chu, X., Moiseenko, A., El Agha, E. (2019). Metformin induces lipogenic differentiation in myofibroblasts to reverse lung fibrosis. Nat Commun, 10(1), 2987. doi:10.1038/s41467-019-10839-0

18. Klee, S., Picart-Armada, S., Wenger, K., Birk, G., Quast, K., Veyel, D., Kastle, M. (2023). Transcriptomic and proteomic profiling of young and old mice in the bleomycin model reveals high similarity. Am J Physiol Lung Cell Mol Physiol, 324(3), L245–L258. doi:10.1152/ajplung.00253.2021

19. Korsunsky, I., Millard, N., Fan, J., Slowikowski, K., Zhang, F., Wei, K., Raychaudhuri, S. (2019). Fast, sensitive and accurate integration of single-cell data with Harmony. Nat Methods, 16(12), 1289–1296. doi:10.1038/s41592-019-0619-0

20. Liang, J., Huang, G., Liu, X., Liu, N., Taghavifar, F., Dai, K., Jiang, D. (2023). Reciprocal interactions between alveolar progenitor dysfunction and aging promote lung fibrosis. Elife, 12. doi:10.7554/eLife.85415

21. McQualter, J. L., McCarty, R. C., Van der Velden, J., O’Donoghue, R. J., Asselin-Labat, M. L., Bozinovski, S., & Bertoncello, I. (2013). TGF-beta signaling in stromal cells acts upstream of FGF-10 to regulate epithelial stem cell growth in the adult lung. Stem Cell Res, 11(3), 1222–1233. doi:10.1016/j.scr.2013.08.007

22. Morse, C., Tabib, T., Sembrat, J., Buschur, K. L., Bittar, H. T., Valenzi, E., Lafyatis, R. (2019). Proliferating SPP1/MERTK-expressing macrophages in idiopathic pulmonary fibrosis. Eur Respir J, 54(2). doi:10.1183/13993003.02441-2018

23. Muniyappa, R., & Gubbi, S. (2020). COVID-19 pandemic, coronaviruses, and diabetes mellitus. Am J Physiol Endocrinol Metab, 318(5), E736–E741. doi:10.1152/ajpendo.00124.2020

24. Natri, H. M., Del Azodi, C. B., Peter, L., Taylor, C. J., Chugh, S., Kendle, R., Banovich, N. E. (2023). Cell type-specific and disease-associated eQTL in the human lung. bioRxiv. doi:10.1101/2023.03.17.533161

25. Nouri-Keshtkar, M., Taghizadeh, S., Farhadi, A., Ezaddoustdar, A., Vesali, S., Hosseini, R., Tahamtani, Y. (2021). Potential Impact of Diabetes and Obesity on Alveolar Type 2 (AT2)-Lipofibroblast (LIF) Interactions After COVID-19 Infection. Front Cell Dev Biol, 9, 676150. doi:10.3389/fcell.2021.676150

26. O’Hare, K. H., & Sheridan, M. N. (1970). Electron microscopic observations on the morphogenesis of the albino rat lung, with special reference to pulmonary epithelial cells. Am J Anat, 127(2), 181–205. doi:10.1002/aja.1001270205

27. Pacheco, E. V., Marega, M., Lingampally, A., Fassy, J., Truchi, M., Goth, K., Rivetti, S. (2023). doi:10.1101/2023.12.22.572972

28. Phipson, B., Sim, C. B., Porrello, E. R., Hewitt, A. W., Powell, J., & Oshlack, A. (2022). propeller: testing for differences in cell type proportions in single cell data. Bioinformatics, 38(20), 4720–4726. doi:10.1093/bioinformatics/btac582

29. Rangarajan, S., Bone, N. B., Zmijewska, A. A., Jiang, S., Park, D. W., Bernard, K., Zmijewski, J. W. (2018). Metformin reverses established lung fibrosis in a bleomycin model. Nat Med, 24(8), 1121–1127. doi:10.1038/s41591-018-0087-6

30. Redente, E. F., Jacobsen, K. M., Solomon, J. J., Lara, A. R., Faubel, S., Keith, R. C., Riches, D. W. (2011). Age and sex dimorphisms contribute to the severity of bleomycin-induced lung injury and fibrosis. Am J Physiol Lung Cell Mol Physiol, 301(4), L510–518. doi:10.1152/ajplung.00122.2011

31. Selman, M., King, T. E., Pardo, A., American Thoracic, S., European Respiratory, S., & American College of Chest, P. (2001). Idiopathic pulmonary fibrosis: prevailing and evolving hypotheses about its pathogenesis and implications for therapy. Ann Intern Med, 134(2), 136-151. doi:10.7326/0003-4819-134-2-200101160-00015

32. Selman, M., & Pardo, A. (2006). Role of epithelial cells in idiopathic pulmonary fibrosis: from innocent targets to serial killers. Proc Am Thorac Soc, 3(4), 364–372. doi:10.1513/pats.200601-003TK

33. Sgalla, G., Iovene, B., Calvello, M., Ori, M., Varone, F., & Richeldi, L. (2018). Idiopathic pulmonary fibrosis: pathogenesis and management. Respir Res, 19(1), 32. doi:10.1186/s12931-018-0730-2

34. Sikkema, L., Ramirez-Suastegui, C., Strobl, D. C., Gillett, T. E., Zappia, L., Madissoon, E., Theis, F. J. (2023). An integrated cell atlas of the lung in health and disease. Nat Med, 29(6), 1563–1577. doi:10.1038/s41591-023-02327-2

35. Street, K., Risso, D., Fletcher, R. B., Das, D., Ngai, J., Yosef, N., Dudoit, S. (2018). Slingshot: cell lineage and pseudotime inference for single-cell transcriptomics. BMC Genomics, 19(1), 477. doi:10.1186/s12864-018-4772-0

36. Taghizadeh, S., Chao, C. M., Guenther, S., Glaser, L., Gersmann, L., Michel, G., Rivetti, S. (2022). FGF10 Triggers De Novo Alveologenesis in a Bronchopulmonary Dysplasia Model: Impact on Resident Mesenchymal Niche Cells. Stem Cells, 40(6), 605–617. doi:10.1093/stmcls/sxac025

37. Taghizadeh, S., Heiner, M., Vazquez-Armendariz, A. I., Wilhelm, J., Herold, S., Chen, C., Bellusci, S. (2021). Characterization in mice of the resident mesenchymal niche maintaining AT2 stem cell proliferation in homeostasis and disease. Stem Cells, 39(10), 1382–1394. doi:10.1002/stem.3423

38. Tsukui, T., Sun, K. H., Wetter, J. B., Wilson-Kanamori, J. R., Hazelwood, L. A., Henderson, N. C., . Sheppard, D. (2020). Collagen-producing lung cell atlas identifies multiple subsets with distinct localization and relevance to fibrosis. Nat Commun, 11(1), 1920. doi:10.1038/s41467-020-15647-5

39. Vaccaro, C., & Brody, J. S. (1978). Ultrastructure of developing alveoli. I. The role of the interstitial fibroblast. Anat Rec, 192(4), 467–479. doi:10.1002/ar.1091920402

40. Van den Berge, K., Roux de Bezieux, H., Street, K., Saelens, W., Cannoodt, R., Saeys, Y., Clement, L. (2020). Trajectory-based differential expression analysis for single-cell sequencing data. Nat Commun, 11(1), 1201. doi:10.1038/s41467-020-14766-3

41. Weckerle, J., Picart-Armada, S., Klee, S., Bretschneider, T., Luippold, A. H., Rist, W., Veyel, D. (2022). Mapping the metabolomic and lipidomic changes in the bleomycin model of pulmonary fibrosis in young and aged mice. Dis Model Mech, 15(1). doi:10.1242/dmm.049105

42. Wu, H., Yu, Y., Huang, H., Hu, Y., Fu, S., Wang, Z., Tang, N. (2020). Progressive Pulmonary Fibrosis Is Caused by Elevated Mechanical Tension on Alveolar Stem Cells. Cell, 180(1), 107–121 e117. doi:10.1016/j.cell.2019.11.027

43. Yan, X., Zhang, J., Pan, L., Wang, P., Xue, H., Zhang, L., Chen, Y. G. (2011). TSC-22 promotes transforming growth factor beta-mediated cardiac myofibroblast differentiation by antagonizing Smad7 activity. Mol Cell Biol, 31(18), 3700–3709. doi:10.1128/MCB.05448-11

44. Yanai, H., Shteinberg, A., Porat, Z., Budovsky, A., Braiman, A., Ziesche, R., & Fraifeld, V. E. (2015). Cellular senescence-like features of lung fibroblasts derived from idiopathic pulmonary fibrosis patients. Aging (Albany NY), 7(9), 664–672. doi:10.18632/aging.100807

45. Zhang, B., Zhou, X., Qiu, Y., Song, Y., Feng, F., Feng, J., Wang, J. (2020). Clinical characteristics of 82 cases of death from COVID-19. PLoS One, 15(7), e0235458. doi:10.1371/journal.pone.0235458

